# Membrane progesterone receptor signaling reverses hyperglycemia and insulin resistance in obese mice

**DOI:** 10.64898/2026.03.27.714733

**Authors:** Nancy Nader, Lubna Zarif, Shereena Sherif, Jawaher Al Hamaq, Dalal al Qahtani, Raphael Courjaret, Fang Yu, Hanan H Abunada, Praveen Babu Vemulapalli, Sunkyu Choi, Frank Schmidt, Khaled Machaca

## Abstract

Progesterone (P4) plays key roles in reproductive and metabolic function and signals through two receptor classes: classical nuclear receptors that regulate gene transcription and membrane progesterone receptors (mPR) that mediate rapid, non-genomic signaling. Whether mPR signaling influences systemic glucose homeostasis remains unclear. Here, we investigated whether mPR activation regulates glucose homeostasis and insulin sensitivity. Using the selective mPR agonist OD02-0, we show that mPR activation enhances glucose uptake in skeletal muscle and hepatocytes, associated with AMP-activated protein kinase (AMPK) activation. In HepG2 cells, mPR activation induces metabolic reprogramming characterized by reduced mitochondrial respiration and increased glycolytic flux. Pharmacological inhibition of AMPK suppresses this effect, indicating that these responses require AMPK activity. In diet-induced obese mice, chronic mPR activation reduces fasting glucose and insulin levels, improves glucose tolerance, and restores glucose-stimulated insulin secretion without detectable toxicity. Integrated proteomic and phosphoproteomic analyses in mouse liver reveal modulation of AMPK signaling and inhibition of mTORC1. Transcriptomic changes were limited, supporting a predominantly non-genomic mode of action. Together, these findings identify mPR signaling as a regulator of glucose homeostasis that engages central energy-sensing pathways to improve metabolic control in obesity.

## Introduction

Progesterone (P4) signaling plays a crucial role in regulating female reproduction, sperm activation, neuronal function, and the immune system (*1–3*). The classical mode of P4 signaling is through nuclear receptors (nPRs), which function as transcription factors to regulate gene expression and elicit cellular responses (*4*). Additionally, P4 activates a rapid transcription-independent signal transduction cascade through membrane progesterone receptors (mPR), which are integral transmembrane proteins. This mode is referred to as nongenomic signaling. mPR belong to the progesterone and adiponectin (AdipoQ) receptor family (PAQRs), that are ubiquitous in eukaryotes, and consist of 11 receptors with 5 being specific to P4: PAQR5 (mPRγ), PAQR6 (mPRδ), PAQR7 (mPRα), PAQR8 (mPRβ) and PAQR9 (mPRɛ) (*5*).

mPR-mediated signaling is recognized as a key regulator of various biological functions, including those in the nervous and cardiovascular systems, female and male reproductive tissues, and in intestinal, immune and cancer cells (*3, 6–9*). Thus, mPRs are emerging as potential therapeutic targets for hypertension, reproductive disorders, and neurological diseases (*10*).

However, the signal transduction cascade downstream of mPR is not well defined. Several papers argue for a role for G-protein signaling and that mPR themselves are GPCRs (*11*). However, those findings are somewhat controversial because of the topology of mPRs (*12*), which is opposite to that of GPCRs (N-terminal intracellular), as well as functional findings. Our recent study using *Xenopus* oocyte maturation offers insights into these discrepancies as it shows that mPRβ activation produces lipid second messengers that are GCPR ligands (*13*). P4 triggers frog oocyte maturation through activation of mPRβ (PAQR8), which requires clathrin-dependent endocytosis and the adaptor protein APPL1, as well as the kinase AKT2 (*12, 14*). APPL1 acts downstream of both insulin and adiponectin signaling, which helps protect against obesity-related metabolic and cardiovascular disorders (*15, 16*). Additionally, AKT2 is a key player in insulin signaling as it promotes glucose uptake and inhibits gluconeogenesis in insulin-sensitive tissues like muscle, liver, and adipose (*17–19*). APPL1 also activates the energy sensor AMPK, which plays a key role in energy metabolism and is the main hepatic target of metformin, one of the primary treatments for type diabetes (T2DM) (*20, 21*). The ability of mPR to regulate AKT2 via APPL1 and potentially AMPK suggests a role in regulating glucose homeostasis. Moreover, mPR were found to boost the secretion of glucagon-like peptide-1 (GLP-1), a key therapeutic target for diabetes and obesity (*22*), and improve glucose tolerance independently of nPRs (*9*). Notably, mPR’s glucoregulatory effect persists in mice lacking GLP-1 and GIP receptors (*9*), suggesting that mPR activate alternative pathways, which do not involve enteroendocrine gut cells, to modulate glucose homeostasis. Together, these observations suggest that mPR signaling may intersect with core metabolic regulatory networks.

In the present study, we aimed to determine whether mPR activation regulates glucose homeostasis at both the whole-animal and cellular levels, whether it can correct hyperglycaemia and insulin resistance in obesity, and to define the underlying signaling mechanisms.

## Methods

### Lead contact and materials availability

Further information and requests for resources and reagents should be directed to the lead contact, Nancy Nader. All biological materials used in this study are available from the lead contact upon reasonable request or from commercial sources. Antibodies, siRNA, primers, chemicals, and kits are listed in Supplemental Table 5. Antibodies dilutions were prepared according to the manufacturer’s recommendations.

### mPR agonist

The selective membrane progesterone receptor (mPR) agonist Org OD 02-0 (OD 02-0; Axon Medchem, Axon 2085) was used for all in vitro and in vivo studies.

### Cell lines and culture conditions

Human HepG2 hepatoma (ATCC HB-8065) and mouse C2C12 myoblasts (ATCC CRL-1772) were obtained from ATCC and cultured according to manufacturer recommendations. C2C12 cells were differentiated into myotubes by culturing in DMEM supplemented with 2% horse serum for 3–5 days. GLUTag cells were provided by Dr. Daniel Drucker (Mt. Sinai Hospital, University of Toronto) and cultured in high-glucose DMEM supplemented with 10% FBS and antibiotics. An APPL1 knockout C2C12 cell line was generated using CRISPR/Cas9 ribonucleoprotein (RNP) technology (GenScript). Cells transfected with non-targeting sgRNA served as controls. Knockout efficiency was confirmed by Sanger sequencing and Western blotting. All cell lines used in this study were routinely tested for mycoplasma contamination and were confirmed to be mycoplasma-free.

### Assessment of mPR expression

Total RNA was extracted using the RNeasy Kit (Qiagen) and reverse-transcribed according to the manufacturer’s instructions. Quantitative real-time PCR was performed using GAPDH as an internal control. Total protein lysates were prepared in RIPA buffer supplemented with protease and phosphatase inhibitors and analyzed by Western blotting where applicable.

### Signaling studies in HepG2 and C2C12 cells

HEPG2 cells were maintained in growth medium consisting of DMEM supplemented with 20% heat-inactivated fetal bovine serum (HI-FBS) and 1% penicillin-streptomycin (Pen/Strep). C2C12 cells were cultured in growth medium (DMEM + 10% FBS + 1% Pen/Strep) for the first 24 hours, followed by differentiation in DMEM supplemented with 2% horse serum and 1% Pen/Strep for 3–5 days, until the formation of multinucleated myotubes was observed. After Overnight starvation, HepG2 and C2C12 myotubes were treated with ethanol (0.1%, as vehicle control) and OD 02-0 (10^-7^/10^-6^M) for 5, 10, 20 and 30 min, and Insulin for 30 min (10^-7^M). C212C were also treated with OD 02-0 for 10 or 30 min in the presence or absence of RU485 (an inhibitor for nuclear progesterone receptors), or Insulin (10^-7^M) (as indicated in the figures). Cells were lysed in RIPA buffer containing protease and phosphatase inhibitors. Equal amounts of protein (30 μg) were resolved by SDS-PAGE and transferred to PVDF membranes. Membranes were incubated with primary antibodies against phosphorylated and total AMPK, AKT2, GSK3β, and eIF4EBP1, followed by HRP- or infrared-conjugated secondary antibodies. Signal detection was performed using ECL or LI-COR Odyssey imaging. Phospho- and total protein levels were normalized to β-actin or tubulin as loading controls. The resulting ratios were further normalized to each other.

### GLUT4 translocation by immunofluorescence and TIRF microscopy

Differentiated C2C12 myotubes were serum-starved and treated with vehicle, insulin (10 nM), and/or OD 02-0 for 30 min. Cells were fixed in 4% paraformaldehyde, permeabilized, blocked, and incubated with anti-GLUT4 primary antibodies followed by Alexa Fluor–conjugated secondary antibodies. Nuclei were stained with Hoechst 33342. Confocal and total internal reflection fluorescence (TIRF) imaging were performed using a Zeiss microscope. GLUT4 plasma membrane localization was quantified by normalizing TIRF fluorescence to epifluorescence intensity.

### Measurement of glucose uptake by FRET imaging

C2C12 myotubes and HepG2 cells were infected with adenovirus encoding the glucose FRET sensor FLII¹²Pglu-700μδ6 at a multiplicity of infection of 100. To construct the adenovirus expressing glucose FRET sensor FLII12Pglu-700μδ6, FLII12Pglu-700μδ6 fragment from pcDNA3.1 FLII12Pglu-700uDelta6 (*23*) was inserted into BglII/HindIII of pShuttle-CMV (*24*). After treatment with vehicle, insulin, OD 02-0, or insulin plus OD 02-0 for 30 min, cells were stimulated with 1 mM glucose (1 mM). Live-cell imaging was performed using a Zeiss LSM880 confocal microscope (excitation 458 nm; emission 482 nm and 570 nm). YFP/CFP ratios were calculated using ImageJ and normalized to baseline prior to glucose addition.

### Mitochondrial and glycolytic stress tests

Mitochondrial respiration and glycolysis were assessed using Seahorse XF Mito Stress and Glycolysis Stress Test Kits (Agilent). HepG2 cells were seeded in XF96 plates, serum-starved for 2 hours, and treated with vehicle or OD 02-0 prior to analysis. Oxygen consumption rate (OCR) was measured following sequential injections of oligomycin, FCCP, and rotenone/antimycin A. Extracellular acidification rate (ECAR) was measured following sequential injections of glucose, oligomycin, and 2-deoxyglucose. OCR and ECAR values were normalized to total cellular DNA content after Hoescht staining and imaging on the ImageXpress Micro XLS system.

### GLP-1 secretion assays

GLUTag cells were transfected with APPL1 siRNA or non-targeting control siRNA using Lipofectamine RNAiMAX. Forty-eight hours post-transfection, cells were treated with progesterone-BSA (200 nM) for 30 min. Culture media were collected for measurement of active GLP-1 using ELISA and normalized to total cellular protein. Cell lysates were analyzed by Western blotting to confirm APPL1 knockdown.

### Mouse models of diet-induced obesity and treatment

Male C57BL/6J mice were obtained from Jackson Laboratory. All animal experiments were approved by the Weill Cornell Medicine-Qatar Institutional Animal Care and Use Committee. Mice were housed in a specific pathogen-free (SPF) facility under controlled environmental conditions, including a 12 h light–dark cycle, ambient temperature of 22 ± 2°C, and relative humidity of 50–60%. Animals were single housed in ventilated cages with standard bedding and environmental enrichment. Mice had ad libitum access to food and water throughout the study. Health status was monitored regularly by trained staff, and all procedures were conducted in accordance with institutional animal care guidelines. Ten-week-old male C57BL/6 mice were fed either standard chow (10% kcal fat) or high-fat diet (60% kcal fat). Body composition was monitored by TD-NMR, and fasting glucose was measured by glucometer. After confirmation of insulin resistance, mice were randomized into three groups: chow-fed controls, HFD mice receiving vehicle, and HFD mice receiving OD 02-0 (4 mg/kg/day) by oral gavage for 28 days. Fasting glucose, insulin levels, glucose tolerance tests (GTT), and glucose-stimulated insulin secretion (GSIS) were assessed. Liver and soleus muscle were collected at sacrifice.

### Glucose tolerance tests and insulin measurements

Mice were fasted overnight prior to GTT. Glucose (2 g/kg) was administered intraperitoneally, and blood glucose was measured at 0, 15, 30, 60, and 120 min. Plasma insulin was measured before and 30–50 min after glucose administration using ultrasensitive mouse insulin ELISA.

### Toxicology and toxicokinetic studies

The objectives of the study were to evaluate the potential toxicity of Org OD02-0 when administered once daily at dose level of 4, 10, and 20 mg/kg by oral gavage to CD-1 mice (22 to 24 weeks of age) for a period of 28 days. Mice were assigned to 7 main (6/sex/group) and 3 toxicokinetic (TK) (9/sex/group) groups and were given once daily oral dose of the vehicle (10% (w/v) hydroxypropyl-beta cyclodextrin (HPBCD) in purified water) and the test item at the dose level of 4, 10 and 20 mg/kg for 28 consecutive days. Toxicity was assessed based on clinical signs, daily body weights, organ weights, clinical, gross and anatomic pathology. Hematology, clinical chemistry, gross pathology and organ weight measurement were performed in all surviving animals of main groups on Day 29. On Day 29 post dose, all surviving animals were euthanized by exsanguination under deep isoflurane anesthesia and animals were subjected to gross necropsy. Histopathology examination was performed on all preserved organs of the vehicle control and high dose groups. Gross changes from all the animals were processed and evaluated microscopically. Histopathology was extended to lower doses (G3 and G4) for thymus in female mice based on observation of test item-related findings at the high dose group, in female mice. The tissues meant for microscopic examination were processed and embedded in paraffin, sectioned, stained with hematoxylin and eosin, and subjected to histopathological examination. Toxicokinetic parameters in plasma were measured. During the in-life phase of the study, blood samples were collected on Day 1 and Day 28 at 30 min, 1 hour, 2-hour, 4-hour, 8 hour and 24 hours post dose by retro orbital plexus from three mice/group/time-point in sparse method of the TK group for bioanalysis. Bioanalysis of plasma samples were conducted at the bioanalytical laboratory, Syngene International Limited and bioanalysis data were used for toxicokinetic calculation. Briefly, toxicokinetic evaluation for plasma concentrations of Org OD 02-0 (Day 1 and Day 28) was performed using validated Phoenix^®^ WinNonlin^®^ software (8.2) (Certara, USA). Noncompartmental toxicokinetic analysis was adopted to obtain the pharmacokinetic parameters from the concentration profiles of Org OD 02-0. Parameters analyzed using extravascular model included “Area under the (concentration time) Curve” (AUC0-24h), peak plasma concentration (Cmax), time to peak plasma concentration (Tmax), Clast and Tlast. When deemed appropriate, the AUC was extrapolated to the infinity (AUC0-inf_obs) by adding to the AUC0-last the ratio of Clast/λz, where λz is the terminal rate constant. Apparent terminal half-life was determined, whenever possible, by linear regression analysis of a minimum of three concentrations after Cmax that are on the terminal phase of the mean concentration–time curve.

### Glycogen measurements in liver

Following fixation, tissues were briefly rinsed in 1× PBS and, if necessary, stored temporarily in 80% ethanol. Dehydration was carried out in three changes of 95% ethanol for 15–20 minutes each, followed by two changes of absolute ethanol for 45 minutes each. Tissues were then cleared in a 1:1 solution of absolute ethanol and xylene for 15 minutes, followed by two changes of pure xylene for 15 minutes each. Infiltration was performed by incubating tissues in molten paraffin at 55 °C overnight. The next day, tissues were embedded in paraffin blocks for microtomy. Paraffin-embedded sections were then cut and mounted onto glass slides, then deparaffinized and rehydrated through a descending alcohol series. For diastase (α-amylase) digestion, duplicate slides were prepared: one designated for digestion and the other for Periodic Acid–Schiff (PAS) staining only. A working α-amylase solution was freshly prepared by dissolving 0.2 g of α-amylase in 40 mL of deionized water. Slides designated for digestion were incubated in the α-amylase solution at 37 °C for 1 hour, followed by a 5-minute wash in running tap water. PAS staining was then performed on both digested and control slides. Slides were oxidized in 1% periodic acid for 5 minutes and rinsed in distilled water. They were then incubated in Schiff’s reagent for 15 minutes, followed by a 10-minute wash in running tap water to develop the magenta coloration. Slides were counterstained with Gill’s hematoxylin for 1 minute, rinsed briefly in water, dehydrated in ascending alcohol grades, cleared in xylene, and mounted with coverslips. This protocol allows for the differentiation of glycogen from other PAS-positive substances by comparing digested and undigested sections. Slides were mounted using Permount mounting medium (Fisher Scientific, SP-15) and allowed to cure for 24 hours before imaging. Slides were then imaged using a Zeiss Axioscan.Z1 slide scanner at 20× magnification. Image acquisition and processing were conducted using Zeiss ZEN software, employing automatic white balance and autofocus functions according to the default scanning protocol optimized for H&E-stained tissue sections. Automated image analysis was performed using Cell Profiler (*25*), the detailed pipeline is available upon request. Briefly, watersheding was performed to separate touching elements and clear objects (lipid droplets) detected and measured for individual size and density. The same objects were then used as a mask and the regions excluded from the PAS staining analysis. Color unmixing was performed using the built-in “PAS” setting and image intensity measured.

### Transcriptomic and proteomic analyses

At the end of the oral gavage experiment, mice were sacrificed, and insulin-sensitive tissues were collected and flash frozen in liquid nitrogen.

#### Transcriptomic

For total RNA preparation, flash frozen Livers were homogenized in 1 ml TRIZOL. The homogenized samples were incubated for 5 minutes at room temperature (RT) then 0.2 ml of chloroform were added. Tubes were shaken vigorously by hand for 15 seconds and incubated at RT for 2 to 3 minutes. Samples were centrifuged 12,000xg for 15 minutes at 2 to 8°C. Following centrifugation, the colorless upper aqueous phase was collected, and one volume of 70% ethanol was added. From this step and onwards, the RNeasy Kit from Qiagen was used to finalize the total RNA preparation. For total RNA preparation from the muscle’s tissues, the RNeasy® Fibrous Tissue kit from Qiagen was used. RNA concentration was measured using a nanodrop and the quality of the RNA was checked by measuring the wavelength absorbance ratios at 260 nm over 280 nm, as well as 260 nm over 230 nm.RNA-sequencing (RNA-seq) of total RNA was conducted at the WCMQ Genomics Core Facility. Briefly, Following RNA extraction, 400ng of total RNA was depleted of ribosomal RNA using RiboCop rRNA Depletion Kit for Human/Mouse/Rat. The rRNA-depleted RNA was used to generate strand-specific ∼ 550bp libraries with CORALL mRNA-Seq V2 Library Prep Kits according to the manufacturer’s protocol. Library quality and quantity were analyzed with the Bioanalyzer 2100 (Agilent, 5067-4626) on High Sensitivity DNA chips. The libraries were then pooled in equimolar ratios and sequenced on Illumina NovaSeq 6000 S1 v1.5, paired-end 100bp run.

### Total and phosho-proteomic

For total proteins preparation, around 50 mg of the flash frozen Livers and muscles were homogenized in RIPA buffer supplemented with proteases and phosphatases inhibitors. Protein concentrations were measured using a BCA protein assay kit.

### Sample preparation for LC/MS/MS analysis

Protein digestion was performed using the PreOmics iST 96x kit (PreOmics GmbH, Martinsried, Germany). Briefly, protein samples were transferred into a 96-well plate using the Agilent Bravo automated liquid handler. Each sample was then brought to a final volume of 50 µL with LYSE buffer using the Bravo system to facilitate protein solubilization, reduction, and alkylation. Samples were incubated at 95°C for 10 minutes.

For digestion, 50 µL of resuspended DIGEST buffer was added to each well, and the plate was incubated at 37°C for 3 hours. The digestion reaction was quenched by adding 100 µL of STOP buffer. Peptides were then transferred to cartridges for cleanup. Peptide cleanup was performed by sequentially washing samples with 200 µL of WASH buffer 1 followed by 200 µL of WASH buffer 2. Peptides were eluted twice with 100 µL of ELUTE buffer to ensure complete recovery. The eluates were dried using a vacuum concentrator and reconstituted in 200 µL of loading buffer (80% acetonitrile, 0.1% TFA) for subsequent phosphopeptide enrichment.

### Phospho-peptide enrichment

Phosphopeptide enrichment was performed using the AssayMAP Bravo system with Fe(III)-NTA cartridges. The cartridges were first primed with 250 µL of priming buffer (50% acetonitrile, 0.1% TFA) at a flow rate of 300 µL/min, then equilibrated with 200 µL of loading buffer (80% acetonitrile, 0.1% TFA) at 10 µL/min.

Peptide samples were loaded slowly onto the cartridges at 1.0 µL/min to ensure optimal binding of Phosphopeptide. To remove any non-specific binders, the cartridges were washed with 250 µL of wash buffer (80% acetonitrile, 0.1% TFA) at 10 µL/min. Phosphopeptide were then eluted using 25 µL of 2% ammonium hydroxide at 2 µL/min and immediately neutralized in 25 µL of 10% formic acid. The eluates were dried using a vacuum concentrator, after dried, stored at −80°C until they were ready for LC-MS/MS analysis.

### MS acquisition

Peptide samples were analyzed on a Q-exactive Explorer 480 (Thermo Scientific) equipped with a nano-flow chromatographic system (Vanquish Neo UHPLC system, Thermo Scientific) following DDA method. Each sample was injected by auto sampler and trapped into Acclaim PepMap 100 Nano-Trap Column (particle size 5 μm, 100Å, I.D.100, length 2 cm, Thermo Scientific), and then separated with EASY-spray PepMap Neo UHPLC C18 column (particle size 2 μm, 100Å, I.D.75 μm, length 50 cm, Thermo Scientific) by a 180 min gradient from buffer A (0.1% formic acid in distilled water) to buffer B (80% ACN, 0.1% formic acid) ) in 45°C column temperature. Eluted peptides were ionized and introduced into the Orbitrap Exploris 480 mass spectrometer (Thermo Scientific) by EASY-Spray Source (Thermo Scientific) with 2.3 kV spray voltage. Full scan was acquired with 400-1650 m/z scan range at 120 k resolution, normalized automatic gain control (AGC) target 300%, 1 micro scan, and auto maximum injection time mode. For MS/MS scan, data dependent analysis was applied with 30 k resolution, 60 second dynamic exclusion, 2 isolation window (m/z), custom AGC target mode, 250% normalized AGC target, auto maximum injection time mode, 1 microscans. For fragmentation of the precursor ions, 28% HCD collision energy was used. Peptide samples were also analyzed in a TimsTOF HT connected to the Evosep One LC system using the 30SPD 44-min standard gradient and EvoSep EV1109 Performance columns (8 cm by 150 μm, 1.5-μm C18 packing). For total proteome analysis, tryptic peptides were loaded onto EvoTip Pure sample tips using Bravo. Peptide samples were analyzed using data-independent acquisition (DIA) in the parallel accumulation–serial fragmentation (PASEF) diaPASEF mode. The diaPASEF method used 32 MS/MS windows per cycle, consisting of one MS1 scan followed by 16 MS/MS scans, and covered the mass range of 400 to 1201 m/z with a 1 m/z overlap between windows to ensure comprehensive coverage. The acquisition spanned the full ion mobility range from 0.60 to 1.60 1/K₀, and each cycle took approximately 1.8 seconds. Collision energy was dynamically adjusted based on ion mobility, ranging from 20 eV at 0.85 1/K₀ to 59 eV at 1.30 1/K₀, to optimize fragmentation efficiency across the range of peptides.

### Data analysis

For DDA analysis, Raw data were processed for protein identification and quantification by MaxQuant (version 1.6.17.0) using the Mus musculus UniprotKB database (release 2024_10, 54,727 protein entries). For modification search, carbamidomethylation (+57.021 Da) of cysteine was selected as a fixed modification, coupled with N-terminal acetylation (+42.011 Da), oxidation (+15.995 Da) of methionine, and phosphorylation (+79.966 Da) of serine and threonine, and Tyrosine were selected as variable modifications. The search tolerance parameters were 20 ppm for first search mass accuracy tolerance, 4.5 ppm for main search mass accuracy, and 20 ppm for FTMS MS/MS tolerance search. The DIA-MS raw data files were processed for protein identification and quantification by Spectronaut 19 software (Biognosys) with library-free approach (direct DIA) mode using the Mus musculus UniprotKB database (release 2024_10, 54,727 protein entries). For modification search, carbamidomethylation (+57.021 Da) of cysteine was selected as a fixed modification, coupled with N-terminal acetylation (+42.011 Da) and oxidation (+15.995 Da) of methionine were selected as variable modifications. For phosphorylation analysis, differential valuable modification of +79.966 Da on serine and threonine, and Tyrosine was also applied. For other search parameters, 5 to 52 amino acids for the peptide length, 20 ppm tolerance for MS and MS/MS search, run wise imputing for imputation strategy, and sum peptide quantity for major quantity, single hit exclusion, and label-free quantification (LFQ) method were applied. In the results, proteins that were quantified with probability value (p-value) less than 0.05 and fold ratio more than 1.5-fold were considered for further analysis. Statistical analysis of total proteome data was performed by SpectroPipeR (*26*), an R package designed to streamline data analysis tasks and provide a comprehensive, standardized pipeline for Spectronaut DIA-MS data. For the phosphoproteomics analyses peptides that are significantly (p<0.05, t-test) up or down regulated in the OD 02-0 compared to the HFD vehicle treatment based on phosphopeptide intensity following enrichment were selected. For the phosphopeptide run total protein intensity was also quantified, and significantly enriched proteins (up or down) were selected. Phosphopeptides that were enriched in the same direction as total proteins were eliminated could be explained by changes in protein levels rather than specific phosphorylation events. Furthermore, only peptides enriched by 2-fold or more (up or down) were selected.

### Statistical studies

Data are presented as mean ± SEM. Each set of experiments was repeated at least 3 times. Groups were compared using the Prism 10 software (GraphPad) using the statistical tests indicated in the figure legend. Statistical significance is indicated by p values (ns: not significant; *** p ≤ 0.001; ** p ≤ 0.01; * p ≤ 0.05).

## Results

### mPR signaling enhances glucose uptake in skeletal muscle and in hepatocytes

The discovery that mPR signals through APPL1 and AKT2 in frog oocytes suggested a role for mPR signaling in mammalian glucose regulation, since both proteins are involved in glycemic control (Fig. 1A). To test this possibility, we examined whether activating mPR engages glucose-regulatory pathways in skeletal muscle, a major site of glucose uptake. We used the selective mPR agonist OD02-0, which does not activate nuclear PRs (*27*).

**Figure 1.**
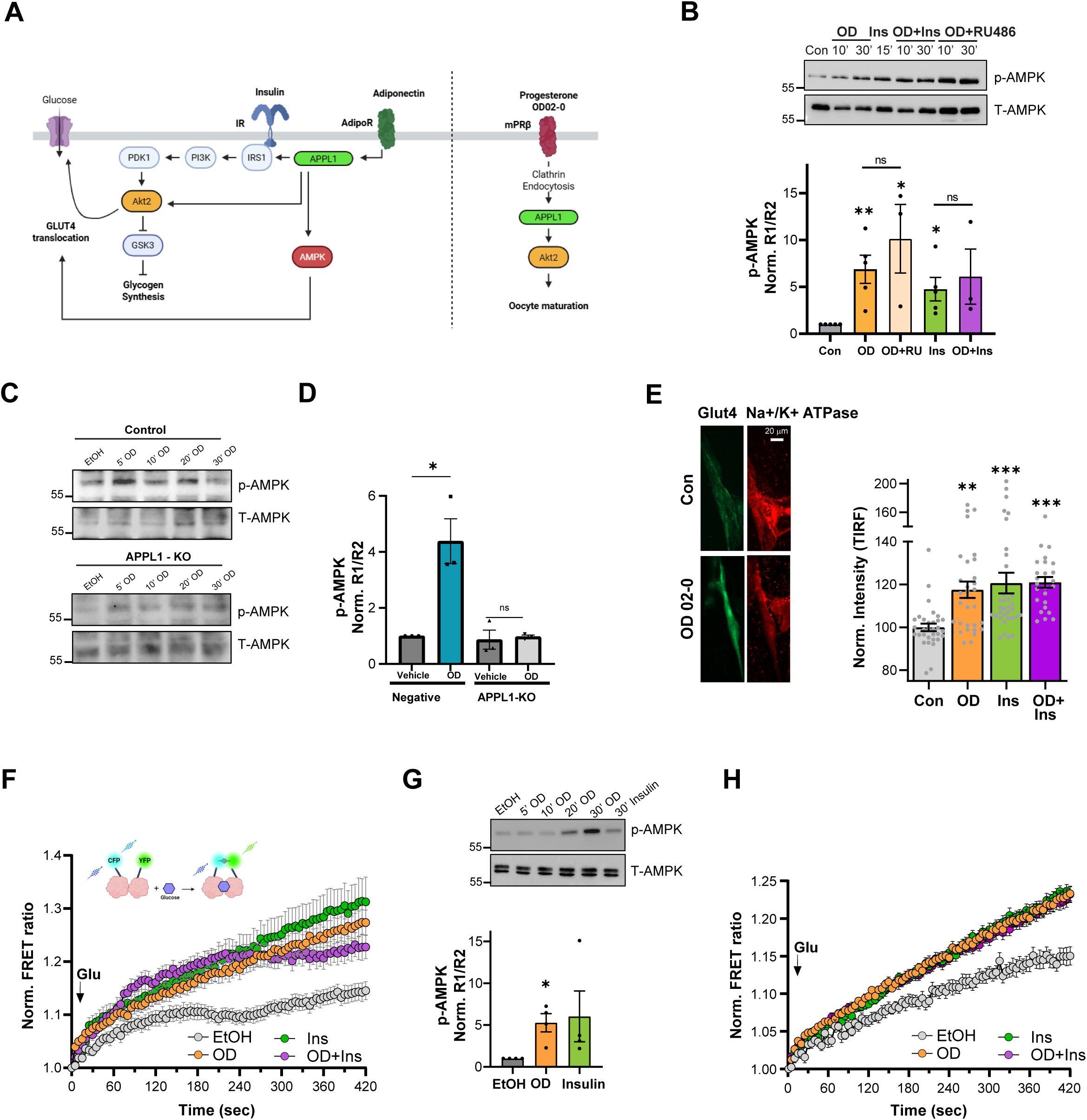
Activation of mPR by OD 02-0 activates AMPK and induces glucose uptake in C2C12 and HepG2 cells. **(A)** Schematic representation of insulin/adiponectin and mPRβ signaling pathways in mammalian muscles cells and *Xenopus laevis* oocytes, respectively. **(B)** Representative immunoblots (*upper*) and quantification (*lower*) of AMPK phosphorylation in differentiated C2C12 myotubes treated with vehicle (EtOH), OD 02-0 (10- or 30-min OD), insulin (15 min), OD + insulin, or OD + RU486. Densitometric quantification for p-AMPK was performed using ImageJ. For each sample, phospho-AMPK/loading control (R1) and total-AMPK/loading control (R2) ratios were calculated, and phosphorylation was calculated as R1/R2. Only 30 min treatment with OD alone or +/- Insulin or RU482 were included in the graph. Final values were normalized to the vehicle (EtOH) (Norm. R1/R2). β-actin served. as loading control (see Supp. Fig. 1E). Data are mean ± SEM, n = 3-5 biological replicates (unpaired two-tailed *t* test). **(C)** Representative immunoblots of phospho- and total AMPK in control and APPL1 knockout (APPL1-KO) C2C12 myotubes treated with EtOH or OD 02-0 (see Supp. Fig. 2C). **(D)** Quantification of AMPK phosphorylation in control and APPL1-KO C2C12 myotubes. For each experiment, the OD time point yielding maximal AMPK phosphorylation was used. Data are mean ± SEM, n = 3–4 independent experiments (unpaired two-tailed *t* test). **(E)** TIRF imaging of endogenous GLUT4 membrane localization in C2C12 myotubes following OD treatment. *Left*, representative images. *Right*, quantification normalized to vehicle. Na⁺/K⁺-ATPase marks the plasma membrane. Scale bar, 20 μm. Data represent 25–32 cells per condition from two independent experiments (one-way ANOVA). **(F)** FLII¹²Pglu-700μδ6 FRET glucose sensor schematic (*upper*) and time course of normalized FRET ratio following glucose addition in C2C12 myotubes pretreated with vehicle, OD, insulin, or OD + insulin for 30 min (*lower*). **(G)** AMPK phosphorylation in HepG2 cells treated with EtOH, OD (5 to 30 min), or insulin. *Upper*, representative immunoblots. *Lower*, quantification using the OD time point yielding maximal AMPK phosphorylation. Tubulin served as loading control (see Supp. Fig. 3G). Data are mean ± SEM, n = 3 independent experiments (unpaired two-tailed *t* test). **(H)** Time course of normalized FRET ratio following glucose addition in HepG2 cells expressing the FLII¹²Pglu-700μδ6 glucose sensor after pretreatment with vehicle, OD, insulin, or OD + insulin. Statistical significance: *p* < 0.05; p < 0.01; *p* < 0.001; ns, not significant.

In skeletal muscle, glucose homeostasis is largely maintained through signals that promote GLUT4 translocation to the plasma membrane to uptake glucose. Insulin drives this process via the IRS-PI3K-Akt pathway (*28, 29*). C2C12 myocytes express two mPRs, mPRα (PAQR7) and mPRβ (PAQR8) (Supp. Fig. S1A). In differentiated C2C12 myotubes (Supp. Fig. S1B), OD02-0 induced only modest AKT2 phosphorylation compared with insulin (Supp. Fig. S1C), and this effect was independent of nuclear P4 receptors (nPRs), as it was not affected by the nPR inhibitor, RU486 (*30*) (Supp. Fig. S1C). Furthermore, no synergy was observed between OD02-0 and insulin in terms of AKT2 activation (Supp. Fig. S1C). Moreover, the downstream AKT2 target GSK3β was phosphorylated (inhibited) to similar levels following OD02-0 or insulin treatment (Supp. Fig. S1D). In addition, OD02-0 activated AMPK (Fig. 1B and Supp. Fig. S1E). Like AKT2, OD02-0–dependent phosphorylation of both GSK3β and AMPK was independent of nPR and did not synergize with insulin (Fig. 1B, Supp. Fig. S1D-E).

As mPR signaling in the oocyte engages APPL1, we were interested in testing the requirement for APPL1 in C2C12 cells. We generated a C2C12 APPL1 knockout (KO) cell line using CRISPR-Cas9 (Supp. Fig. S2A-B). OD02-0-induced phosphorylation of AKT2, AMPK, and GSK was abolished in the knockout cells (Fig. 1C-D, Supp. Fig. S2C-E), demonstrating that mPR signaling in C2C12 cells requires APPL1. Interestingly, APPL1-deficient cells displayed constitutively elevated inhibitory phosphorylation of GSK3 independent of AKT2 (Supp. Fig. S2E). We next asked whether the APPL1 dependency was maintained in another relevant cell type. We tested the murine enteroendocrine GLUTag cells, because it secretes GLP-1 in response to P4 in an mPR-dependent manner (*9*). Treating GLUTag cells with P4-BSA (mPR-selective as it is not permeant to the PM) failed to induce GLP-1 secretion when APPL1 expression was downregulated (Supp. Fig. S2F-G). Together, these findings suggest that APPL1 acts as a conserved adaptor for mPR signaling.

Because both AKT2 and AMPK promote GLUT4 translocation to the plasma membrane (PM), we assessed whether OD02-0 alters GLUT4 trafficking. Indeed, OD02-0 significantly increased GLUT4 PM localization, comparable to insulin, as shown by TIRF imaging (Fig. 1E) and confocal microscopy (Supp. Fig. S3A). OD02-0 and insulin did not act synergistically (Fig. 1E*, right panel*). Consistent with the GLUT4 translocation data, OD02-0 increased glucose uptake, measured using a FRET-based glucose sensor (Supp. Fig. S3B), to a similar extent as insulin (Fig. 1F). As expected, cytochalasin treatment reduced basal glucose uptake, consistent with its known ability to disrupt GLUT4-dependent glucose transport (Supp. Fig. S3C) (*31*).

We next tested mPR signaling in hepatocytes because the liver plays a central role in glucose homeostasis by balancing glucose production, storage, and utilization. Unlike skeletal muscle and adipose tissue, where insulin acutely increases glucose uptake by recruiting GLUT4 to the PM, hepatocytes express GLUT2, a high-capacity bidirectional transporter that constitutively localizes at the cell surface (*32*). Consequently, insulin does not regulate hepatic glucose entry through transporter translocation and abundance at the PM. Instead, it enhances hepatic glucose uptake indirectly by accelerating intracellular glucose phosphorylation and utilization, thereby sustaining the concentration gradient that drives glucose influx (*33, 34*).

We used human HepG2 hepatocytes to determine whether mPR signaling intersects with these metabolic processes. Among the five mPR proteins, HepG2 cells predominantly express mPRβ (PAQR8), and to a lesser extent mPRα and mPRγ (PAQR7 and PAQR5, respectively) (Supp. Fig. S3D). OD02-0 did not activate AKT2 (Supp. Fig. S3E) but induced GSK3 phosphorylation independently of AKT2, suggesting an alternative upstream mechanism (Supp. Fig. S3F). Consistent with our findings in C2C12 cells, OD02-0 triggered robust AMPK activation (Fig. 1G and Supp. Fig. S3G) and glucose uptake (Fig.1H) to similar levels to insulin.

Together, the findings in C2C12 and HepG2 cells show that mPR activation induces AMPK phosphorylation and increases glucose uptake, two key metabolic pathways.

### OD02-0 reverses hyperglycemia and corrects insulin resistance in obese mice

To determine whether these cellular effects translate to whole-body glucose homeostasis, we next tested the effects of OD02-0 in a mouse model of diet-induced hyperglycemia and insulin resistance. Age-matched male mice were placed on chow diet (Group 1), or a high-fat diet (HFD) for 10 weeks (Groups 2 and 3) (Fig. 2A). HFD feeding resulted in progressive weight gain (Supp. Fig. S3H and Fig. 2B) and a significant increase in fat mass (Fig. 2C). It also induced hyperglycemia, as shown by elevated fasting glucose levels (Fig. 2D). After the onset of hyperglycemia, HFD-fed mice received daily oral treatment with either vehicle (HFD-V) or OD02-0 (HFD-OD) for 28 days (Fig. 2A). Although OD02-0 did not alter body weight or fat percentage relative to vehicle-treated HFD mice (Fig. 2B-C), it markedly reversed obesity-associated hyperglycemia (Fig. 2D). Average fasting glucose levels in the HFD-OD group dropped from 204.7 ± 4.15 mg/dl to 165.9 ± 6.79 mg/dl (p < 0.001, n = 16), with 14 of 16 mice showing substantial improvement. Notably, fasting glucose levels in the HFD-OD group were no longer significantly different from lean controls (152.5 ± 3.9 mg/dl, p = 0.53, n = 15) and were significantly lower than HFD-V mice (199.9 ± 8.78 mg/dl, p < 0.001, n = 13) (Fig. 2D).

**Figure 2.**
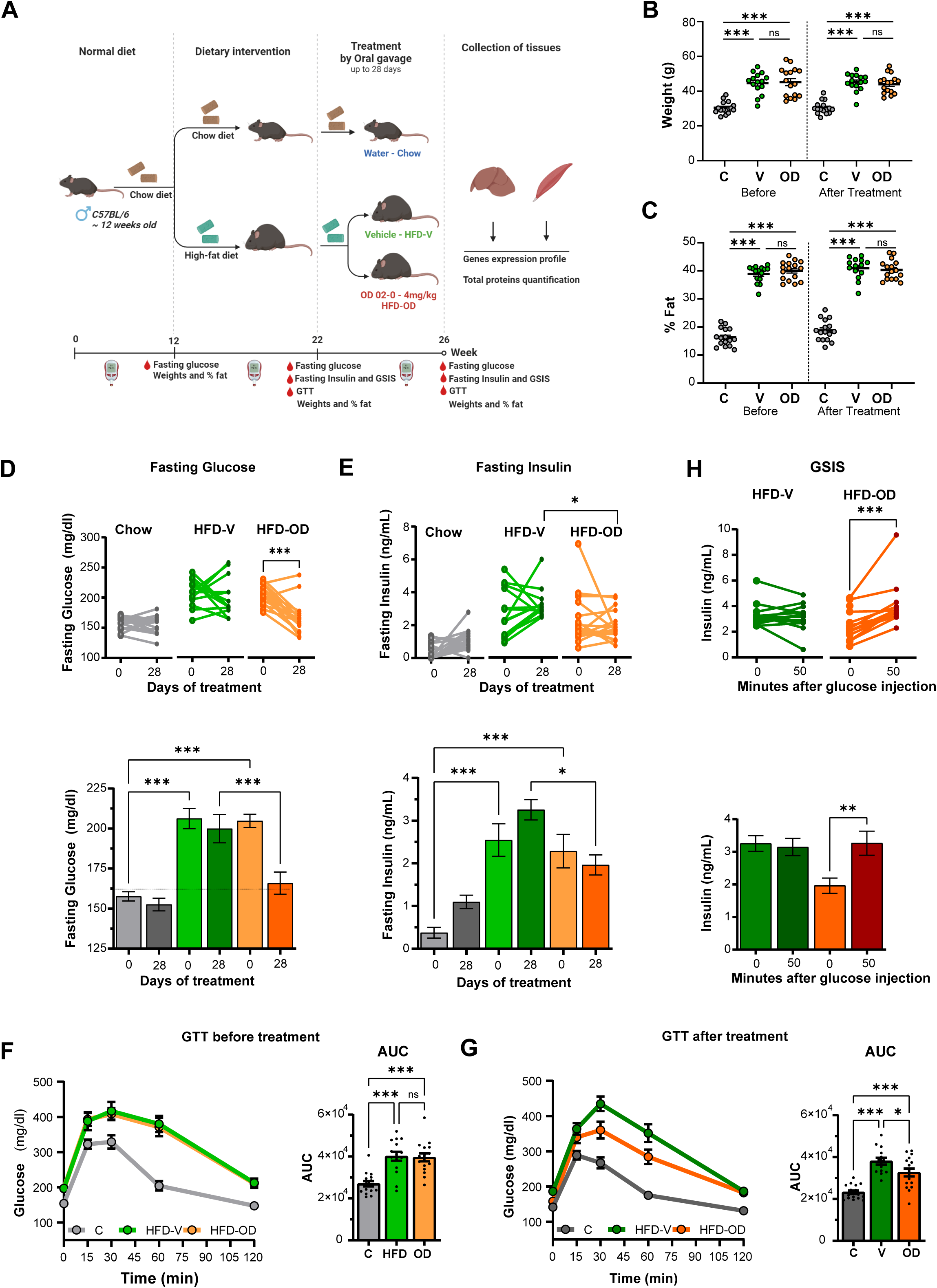
OD 02-0 reverses hyperglycemia and insulin resistance in high-fat diet-induced obese mice. **(A)** Experimental timeline. Male C57BL/6J mice were fed standard chow or high-fat diet (HFD) for 10 weeks, followed by daily oral gavage of vehicle (HFD-V) or OD 02-0 (HFD-OD) for 4 weeks while maintained on diet. **(B, C)** Body weight (B) and fat mass percentage (C) were measured before and after treatment. Data are mean ± SEM, n = 13–16 mice per group from two independent experiments (one-way ANOVA). **(D, E)** Fasting blood glucose (D) and insulin (E) were measured before and 21–28 days after treatment. *Upper panels* show individual values; *lower panels* show group means ± SEM. n = 13–16 mice per group from two independent experiments (one-way ANOVA). **(F, G)** Intraperitoneal glucose tolerance tests were performed before (F) and after (G) treatment. *Left*, glucose levels over time. *Right*, AUC quantification. Data are mean ± SEM, n = 14–16 mice per group from two independent experiments (one-way ANOVA). **(H)** Glucose-stimulated insulin secretion in HFD-fed mice treated with vehicle or OD 02-0. Insulin levels were measured at fasting and 50 min after glucose administration. *Upper panels* show individual values. *Lower panels* show group means ± SEM. n = 14–16 mice per group from two independent experiments (one-way ANOVA). Statistical significance: *p* < 0.05; p < 0.01; *p* < 0.001; ns, not significant.

Moreover, before treatment, fasting insulin levels were significantly elevated in the HFD groups (Group 2: 2.5 ± 0.14 ng/ml, *p* < 0.001, *n* = 15; Group 3: 2.3 ± 0.15 ng/ml, *p* < 0.001, *n* = 16) compared to the lean Chow group (Group 1: 0.43 ± 0.12 ng/ml, *n* = 16), indicating the development of insulin resistance in the obese mice (Fig. 2E). Following treatment with OD02-0, fasting insulin levels in the HFD-OD group significantly decreased (1.9 ± 0.23 ng/ml, *n* = 16, *p* < 0.001) compared to the HFD-V group (3.26 ± 0.23 ng/ml, *n* = 14) (Fig. 2E), with levels sharply decreasing in one animal that showed high fasting insulin before treatment. Whereas over the 28-day period of treatment, fasting insulin levels continued to rise in the HFD-V group, they trended down in the HFD-OD group (Fig. 2E). We further assessed insulin resistance by measuring glucose clearance using the glucose tolerance test (GTT). As expected, before the OD 02-0 treatment, animals on the HFD cleared glucose at a significantly slower rate than their lean counterparts (Fig. 2F), consistent with insulin resistance. OD 02-0 treatment significantly improved glucose clearance compared to the HFD-V group (Fig. 2G). These results show that OD 02-0 treatment effectively reverses insulin resistance.

Finally, we also measured blood insulin levels in response to a glucose injection in fasted animals on the HFD (glucose-stimulated insulin secretion (GSIS); Fig. 2H). Glucose did not alter insulin levels in the HFD-V group, but it was associated with an increase in insulin levels in HFD-OD group (Fig. 2H), showing enhanced pancreatic function in response to a glucose challenge.

### OD02-0 is safe with no associated toxicity or side effects

The above data argue that OD02-0 is a viable therapeutic to reverse hyperglycemia and insulin resistance. During the 28 days of treatment at the 4 mg/Kg dose, we did not observe any overt signs of discomfort or toxicity in the animals. We further evaluated potential OD02-0 toxicity and its pharmacokinetic properties when administered orally at three different doses 4, 10, and 20 mg/kg to male and female CD-1 mice for a period of 28 days. We assessed toxicity based on clinical signs, daily body weights, organ weights, clinical, gross and anatomic pathology. Histopathology examination was performed on organs from vehicle control and 20 mg/kg groups. No mortality or clinical signs were observed in any of the treated groups. Additionally, no significant changes were noted in body weights (Supp. Fig. S4A-B) or food consumption (Supp. Fig. S4C). There were also no test item-related changes in glucose levels or clinical chemistry parameters (Supp. Fig. S4D), as well as the hematological parameters, including red and white blood cell counts (Supp. Fig. S4E-F). At the tissue level no changes in morphology (data not shown) or relative weights were noted in different organs from both sexes (Supp. Fig. S4G). Moreover, and following oral administration of OD02-0 at 4, 10, or 20 mg/kg/day in male and female CD-1 mice, peak plasma concentrations (Cmax) were reached within 0.5 to 1 hour on both Day 1 and Day 28 for both sexes and were mostly cleared within 4 hours (Supp. Table S1 and Supp. Fig. S4H). The half-life of Org OD02-0 varied with sex, dose, and treatment duration. On Day 1, it was quantifiable in females across all doses (3.4–8.1 h) and in males at 10 mg/kg (4.7 h). By Day 28, half-lives were shorter, with detectable values only at 4 mg/kg in males (2.3 h) and at 10 and 20 mg/kg in females (1.6 and 1.2 h), indicating faster clearance with repeated dosing (Supp. Table S1 and Supp. Fig. S4H). Quantifiable concentrations of OD02-0 were detected up to 8-24 hours. Furthermore, no alterations were observed from histopathological studies from an extensive list of over 50 organs and tissues listed in Supp. Table S2. Overall, these findings underscore the therapeutic potential of targeting mPR with OD 02-0 to reverse hyperglycemia and insulin resistance associated with obesity, with no remarkable side effects.

### Omics studies to explore the mechanism of action of mPR in glucose regulation

The beneficial antidiabetic effects of OD02-0 at the whole-animal level are especially compelling given the absence of detectable toxicity, even at doses up to five times higher than the therapeutic regimen. However, despite its broad use as a selective mPR agonist (*35–40*), we cannot exclude the possibility that OD02-0 is metabolized into intermediates capable of activating classical nuclear progesterone receptors (nPRs), a potential concern given the established roles of nPRs in reproductive control and in breast, ovarian, and brain cancers (*41–44*). In addition, mPRs have been reported to influence transcription indirectly through downstream signaling intermediates (*45*). To define the pathways engaged by OD02-0 and to assess whether its effects are primarily non-genomic, we performed integrated transcriptomic, proteomic, and phosphoproteomic analyses in liver and soleus muscle, two major insulin-sensitive tissues that regulate systemic glucose homeostasis.

### Transcriptomic Profiling

RNAseq analyses detected relatively minor differences between chow-fed and HFD animals in muscle (Supp. Fig. 5A). In contrast, in the liver the HFD was associated with significant changes (Supp. Fig. S5B). However, OD02-0 treatment did not substantially alter the hepatic or muscle transcriptomes, as evidenced by the minimal differences between HFD-OD and HFD-V groups (Supp. Fig. S5A-B, *right panels*). This supports the view that OD02-0 acts primarily through non-genomic signaling and does not engage nPR-mediated transcriptional programs.

### Proteomic Remodeling

At the proteomic level, UMAP analysis showed clear separation between the three treatments more so in liver than in muscle (Supp. Fig. S5C), with many changes attributable to diet alone (Supp. Fig. S5D-E). When comparing HFD-V and HFD-OD animals, skeletal muscle proteomes were largely unchanged (Supp. Fig. S5F), indicating that OD02-0 does not induce broad protein remodeling in this tissue. However, because our in vitro data demonstrate that OD02-0 enhances glucose uptake in myotubes, the lack of major proteomic shifts likely reflects functional regulation without large-scale protein turnover. In contrast, OD02-0 induced marked proteomic alterations in the liver (Fig. 3A-B), suggesting that hepatic remodeling may be a key driver of its systemic metabolic effects.

**Figure 3.**
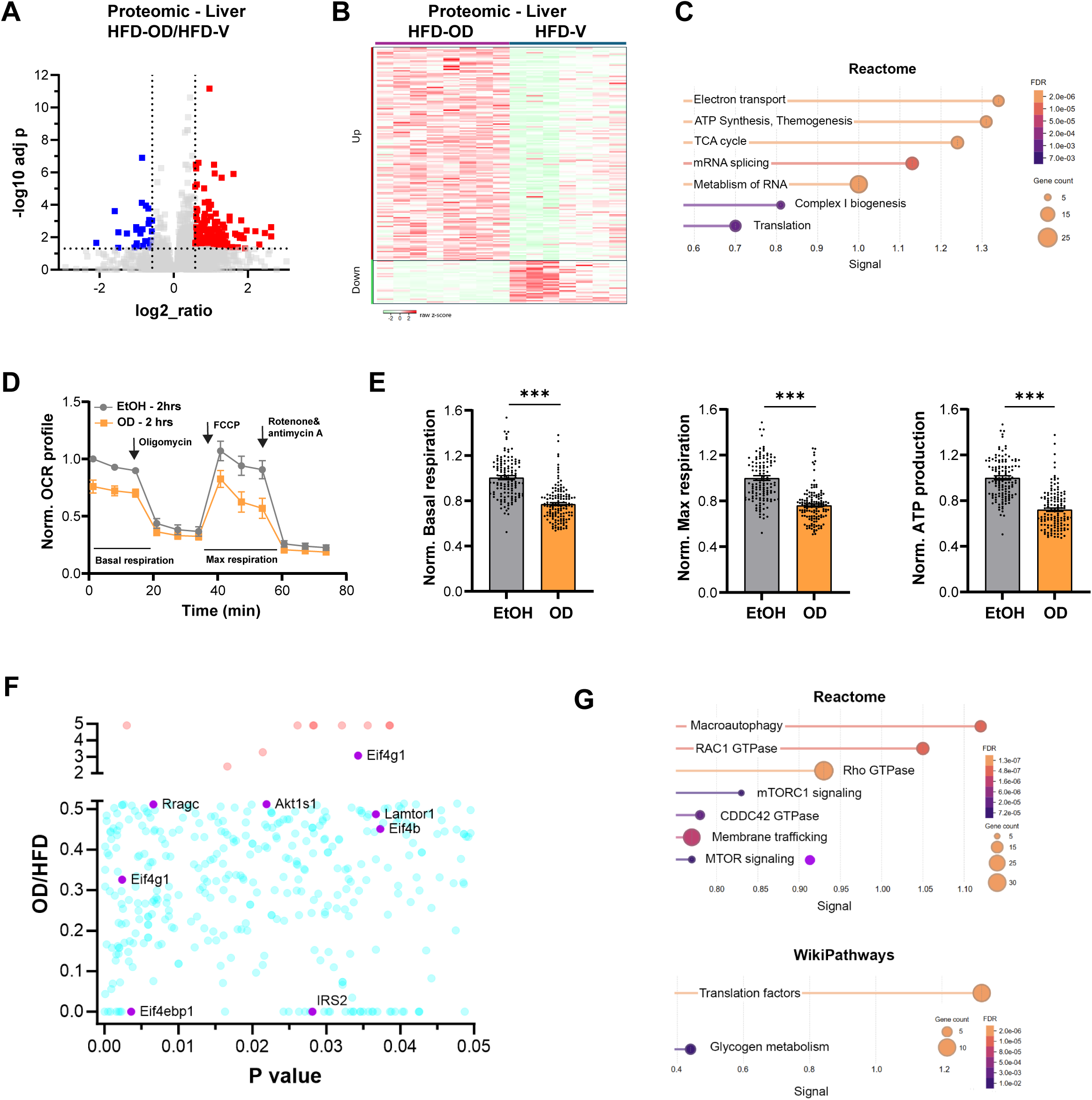
Omics data and regulation of mitochondrial function by OD02-0 in hepatocytes. **(A)** Proteomic analysis of liver tissues from HFD-fed mice treated with OD 02-0 (HFD-OD, n = 8) or vehicle (HFD-V, n = 7). Volcano plot of differentially expressed proteins (fold change > 1.5 or < −1.5; adjusted *p* < 0.05). **(B)** Heatmap of differentially expressed liver proteins from HFD-OD versus HFD-V mice (row Z-score–normalized). **(C)** Pathway enrichment analysis (STRING v12, Reactome) of proteins upregulated in HFD-OD livers. Dot size indicates mapped proteins; color scale represents FDR. **(D, E)** Mitochondrial respiration in HepG2 cells treated with OD 02-0 (10⁻⁶ M) or vehicle for 2 hours, measured by Seahorse XF Mito Stress Test. (D) Oxygen consumption rate (OCR) was measured over time in response to sequential injections of oligomycin, FCCP and rotenone and antimycin A; (E) quantification of basal, maximal, and ATP-linked respiration. Data are mean ± SEM, n = 3 independent experiments (unpaired two-tailed *t* test; *p* < 0.001). **(F)** Scatter plot of phosphoproteins showing phosphorylation ratios (HFD-OD/HFD-V) versus associated *p* values. Hypophosphorylated (blue) and hyperphosphorylated (pink) proteins are indicated. Selected mTOR components are highlighted; IRS2 is shown in red. **(G)** Pathway enrichment analysis (STRING v12, Reactome and WikiPathways) of significantly regulated phosphoproteins. Dot size indicates mapped proteins; color scale represents FDR.

### Mitochondrial Protein Changes and OXPHOS Effects

Pathway enrichment analyses of the subset of proteins reduced in the liver of OD02-0–treated animals revealed suppression of cholesterol biosynthesis (Reactome MMU-191273; FDR = 1.5 × 10⁻⁴; 4/27 proteins), arguing for a potential beneficial effect of OD 02-0 on dyslipidemia, although this was not further explored. In contrast, analyses of proteins increased in response to OD 02-0 in the liver show strong enrichment of pathways related to mitochondrial respiration, ATP production, the TCA cycle, and thermogenesis (Fig. 3C). Among the upregulated hits was NDUFS1, a complex I subunit. Western blotting confirmed increased NDUFS1 protein levels in OD02-0-treated livers relative to vehicle (Supp. Fig. S6A). Consistently, exposing HepG2 cells to OD02-0 also elevated NDUFS1 and the complex III subunit UQCRFS1 (Supp. Fig. S6B).

The increase in proteins involved in mitochondrial respiration in livers from OD02-0 treated animals argued for increased mitochondrial respiration. Surprisingly, treating HepG2 with OD02-0 suppressed oxidative phosphorylation (OXPHOS), reducing both basal and maximal respiration (Fig. 3D-E). This is the opposite phenotype than what would be predicted from the proteomics profiles. However, the decreased respiration would explain the observed activation of AMPK in response to OD02-0, because OD02-0 exposure led to a drop in ATP levels (Fig. 3E, *right panel*), which would activate AMPK as it is classically induced by ATP depletion (*46*). These findings argue that the upregulation of mitochondrial proteins represents a compensatory mechanism to reverse the decrease in OXPHOS induced by OD02-0 and suggest an AMPK-driven remodeling response. In fact, AMPK is well known to coordinate mitochondrial biogenesis and dynamics under metabolic stress (*47*).

### Phosphoproteomics implicate mTORC1 Signaling

Because the observed proteomic shifts occurred without accompanying transcriptional changes, the data point to post-transcriptional regulation, either stabilization of existing proteins or enhanced translation. Translation itself was among the top enriched pathways (Fig. 3C), prompting phosphoproteomic profiling to define upstream regulatory nodes. Phosphoproteomic analyses identified 375 regulated phosphopeptides from 321 proteins in OD02-0–treated livers. Eleven were hyperphosphorylated and the remainder were dephosphorylated with most targets carried two phosphosites, with a minority harboring three or four. (Fig. 3F). Pathway enrichment analysis (STRING v12) revealed significant clustering in macroautophagy, Rho-family GTPases (RAC1, Rho, CDC42), and the mTORC1 pathway (Fig. 3G, Reactome, and Supp. Table S3), as well as translation initiation and glycogen metabolism (Fig. 3G, WikiPathways). Several of the regulated proteins, such as Lamtor1, and Rragc, were shared between the mTORC1 and autophagy pathways. Rho-family GTPases, classically linked to actin remodeling, have also been implicated in insulin secretion and may analogously govern glucose transport in hepatocytes (*48*). A key mTORC1 substrate, Eif4ebp1, was strongly dephosphorylated in OD02-0–treated livers (Fig. 3F), indicating mTORC1 inhibition in vivo. Interestingly, AMPK and mTORC1 are often described as the “yin and yang” of nutrient sensing, with AMPK acting as a negative regulator of mTORC1 activity (*49, 50*). This regulatory axis is consistent with the AMPK activation observed in C2C12 and HepG2 cells following OD treatment (Fig. 1). Therefore, we then validated these observations by Western blot. Livers from OD02-0-treated mice exhibited reduced phosphorylation of Eif4ebp1 and elevated phosphorylation of AMPKα (Supp. Fig. S6C-D), and direct exposure of HepG2 cells to OD02-0 led to time-dependent dephosphorylation of Eif4ebp1 (Supp. Fig. S6E). Since mTORC1 promotes mitochondrial oxidative capacity (*51*), its inhibition might provide a mechanistic explanation for the reduction in OXPHOS observed in HepG2 cells (Fig. 3D-E). The concurrent increase in mitochondrial proteins therefore likely reflects an adaptive response to AMPK activation and suppression of mTORC1 activity.

### Insulin Receptor Substrates and Insulin Sensitivity

Moreover, chronic mTORC1 activation is known to drive insulin resistance by promoting inhibitory phosphorylation and degradation of insulin receptor substrates (IRS) (*52, 53*). Although IRS1 itself was not detected in our phosphoproteomic dataset, we observed significant dephosphorylation of IRS2, the predominant IRS isoform in hepatocytes (Fig. 3F). Specifically, the peptide identified showed reduced phosphorylation at T517, a residue that may play a regulatory role analogous to inhibitory phosphosites described for IRS1. To further explore this, we assessed total IRS1 and IRS2 protein levels in liver lysates. IRS1 was undetectable in HFD-V animals but became discernable in 7 of 8 OD02-0–treated mice (Fig. 4A), suggesting restoration or stabilization of IRS1 expression in response to treatment. Unfortunately, none of the commercially available antibodies yielded a specific IRS2 signal in mouse liver, preventing definitive analysis at the protein level. Nevertheless, the combination of restored IRS1 and reduced IRS2 phosphorylation supports the possibility that mPR activation helps reverse insulin resistance by relieving mTORC1-driven suppression of insulin receptor substrates.

**Figure 4.**
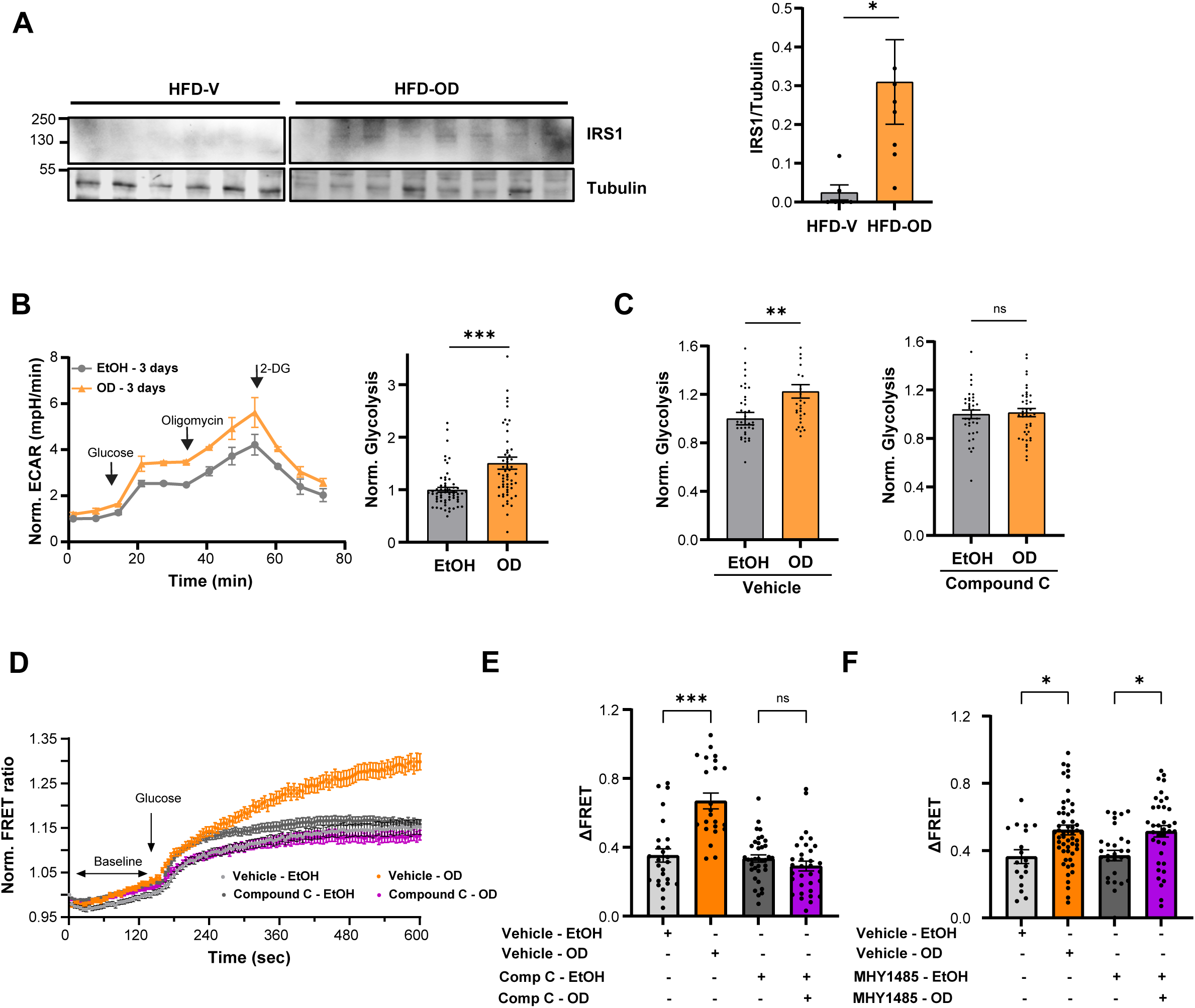
OD 02-0 modulates mTOR and AMPK signaling in mouse liver and HepG2 cells. **(A)** Representative immunoblot (*left*) and quantification of IRS1 protein levels (*right*) in liver tissues from HFD-fed mice treated with vehicle (HFD-V, n = 6) or OD 02-0 (HFD-OD, n = 8). IRS1 was detectable only in HFD-OD samples. IRS1 levels were normalized to tubulin. Data are mean ± SEM. **(B)** Glycolytic function was assessed in HepG2 cells treated with OD02-0 (10⁻⁶ M) or vehicle (EtOH, 0.1%) for 3 days, using the Seahorse XF Glycolysis Stress Test. *Left panel,* extracellular acidification rate (ECAR) was measured over time in response to sequential injections of glucose, oligomycin, and 2-deoxyglucose. *Right panel*; normalized glycolytic rate in HepG2 cells treated with OD 02-0 (10⁻⁶ M) or vehicle (EtOH, 0.1%). Data represent mean ± SEM from 3 independent experiments (unpaired two-tailed *t*-test. **p < 0.01). **(C)** Normalized glycolytic rate to the ethanol condition (EtOH) in HepG2 cells treated with ethanol (*left*) or OD 02-0 (*right*) following pretreatment with the AMPK inhibitor Compound C. Data are mean ± SEM from 31–44 cells per condition (unpaired two-tailed *t* test; p < 0.01). **(D)** Time course of normalized FRET ratio following glucose addition in HepG2 cells expressing the FLII¹²Pglu-700μδ6 glucose sensor after pretreatment with Compound C or vehicle, followed by OD 02-0 or ethanol (EtOH) stimulation. **(E)** Quantification of ΔFRET derived from traces in (D). Data are mean ± SEM from 23–36 cells from two independent experiments (one-way ANOVA; *p* < 0.001; ns). **(F)** Quantification of ΔFRET derived from traces in Sup. Fig. 6G. HepG2 cells expressing the FLII¹²Pglu-700μδ6 glucose sensor were imaged after pretreatment with MHY1485 or vehicle, followed by OD 02-0 or ethanol (EtOH) stimulation. Data represent mean ± SEM from n = 18 to 56 cells from 2 independent experiments (one-way ANOVA. *p < 0.05; ns, not significant).

### OD02-0 Increases Glycolysis through AMPK Activation but independently of mTORC1

Because AMPK stimulates glycolysis in the liver (*54, 55*), we next assessed glycolytic output in response to OD02-0 in HeG2 cells. OD02-0 exposure, for 2 hours or 3 days, significantly increased glycolysis in HepG2 cells, as measured by Seahorse extracellular flux analysis (Supp. Fig. S6F and Fig. 4B). This increased glycolytic flux nicely explains the enhanced glucose uptake in response to OD 02-0 in hepatocytes (Fig. 1H). These results argue that the decrease in OXPHOS observed in response to OD 02-0 results in AMPK activation, which in turn upregulates glycolysis. To test whether this is the case, we blocked AMPK using Compound C, an ATP-competitive inhibitor of AMPK (*56*). Compound C pretreatment prevented both the increase in glycolysis (Fig. 4C) and in glucose uptake (Fig. 4D-E) that are typically induced in response to OD 02-0. These data confirm a requirement for AMPK activation to induce both glycolysis and glucose uptake.

We next asked whether the AMPK dependency of OD02-0-induced glucose uptake requires suppression of mTORC1. We treated HepG2 cells with MHY1485, a lysosomal retention agonist that maintains mTORC1 activity independent of nutrient status (*57*). Despite enforced mTORC1 activation, OD02-0 still increased glucose influx (Supp. Fig. S6G) with ΔFRET values remaining comparable to OD02-0 alone (Fig. 4F). These findings indicate that although AMPK is necessary for OD02-0–driven glucose uptake, this effect does not require downstream inhibition of mTORC1.

### Glycogen and Lipid Metabolism

Because glycogen metabolism emerged as an enriched pathway in the phosphoproteomic analysis (Fig. 3G, and Supp. Table S4), we next examined relevant phosphoproteins. Several enzymes involved in glycogen synthesis and mobilization, including phosphoglucomutase-1 (Pgm1), glycogen synthase (GYS2), and the regulatory subunit of phosphorylase b kinase (Phkb), were dephosphorylated in OD02-0–treated livers (Supp. Table S4). These modifications could impact glycogen metabolism in vivo. Consistent with metabolic dysfunction, mice on HFD displayed markedly reduced hepatic glycogen compared to chow-fed controls (Supp. Fig. S6H). However, glycogen levels were similar between HFD-V and HFD-OD groups (Supp. Fig. S6H, *right panel*), suggesting that in the obese liver, OD02-0 does not further elevate glycogen content, possibly because hepatic glucose is directed toward other metabolic fates under lipotoxic conditions. To determine whether lipid metabolism was affected, we assessed hepatic steatosis, but no differences were detected between OD02-0 and vehicle groups (Supp. Fig. S6I). Thus, the antidiabetic effects of OD02-0 are not attributable to reduced steatosis or changes in liver fat content. Strikingly, despite persistent obesity and fatty liver, OD02-0-treated animals maintained robust glycemic control, indicating that the compound improves insulin sensitivity and glucose homeostasis independently of weight or hepatic lipid remodeling.

## Discussion

Diabetes remains a major and growing global health challenge, encompassing both type 1 diabetes, characterized by autoimmune destruction of pancreatic β-cells, and type 2 diabetes (T2DM), which arises from progressive insulin resistance and impaired glucose homeostasis, leading to significant comorbidities if untreated (*58, 59*). Although lifestyle interventions are foundational, sustained adherence is limited and pharmacological therapies are often required. Despite major therapeutic advances, including GLP-1 receptor agonists, SGLT2 inhibitors, DPP-4 inhibitors, and combination regimens, durable glycemic control is not consistently achieved in patients with T2DM (*60*). First-line oral agents such as metformin, SGLT2 inhibitors, and DPP-4 inhibitors allow only ∼50–60% of patients to reach target glucose levels, and their efficacy declines over time. GLP-1 receptor agonists improve glycemia and support weight loss but are limited by cost, gastrointestinal side effects, and injectable administration. Even bariatric surgery, while highly effective initially, is associated with relapses in a subset of patients (*61–64*). These limitations underscore the need for alternative molecular strategies to restore glucose homeostasis. In this context, exploring novel molecular pathways such as membrane progesterone receptors is appealing.

Here, we demonstrate that activation of membrane progesterone receptors (mPR) improves systemic glucose control in a model of diet-induced obesity. While P4 signaling has traditionally been understood through nuclear receptor-mediated transcriptional regulation, mPR mediate rapid, non-genomic signaling events whose metabolic roles have remained incompletely defined. Our findings support a functional link between mPR activation and key nutrient-sensing pathways that regulate hepatic glucose handling and whole-body insulin sensitivity.

Using integrated cellular, in vivo, and multi-omics approaches, we show that the selective mPR agonist OD02-0 reverses hyperglycemia and improves insulin resistance in obese mice without detectable toxicity. Mechanistically, mPR activation converges on AMPK activation and inhibition of mTORC1 signaling in the liver. Functional studies in hepatocyte models further demonstrate reduced mitochondrial respiration and enhanced glycolytic flux following mPR stimulation, providing a cellular basis for increased glucose uptake and improved glycemic control. In skeletal muscle, mPR activation similarly enhances AMPK signaling and promotes glucose uptake through increased GLUT4 recruitment to the plasma membrane (Fig. 5). Together, these data position mPR signaling upstream of established metabolic nodes central to energy balance and insulin responsiveness.

**Figure 5.**
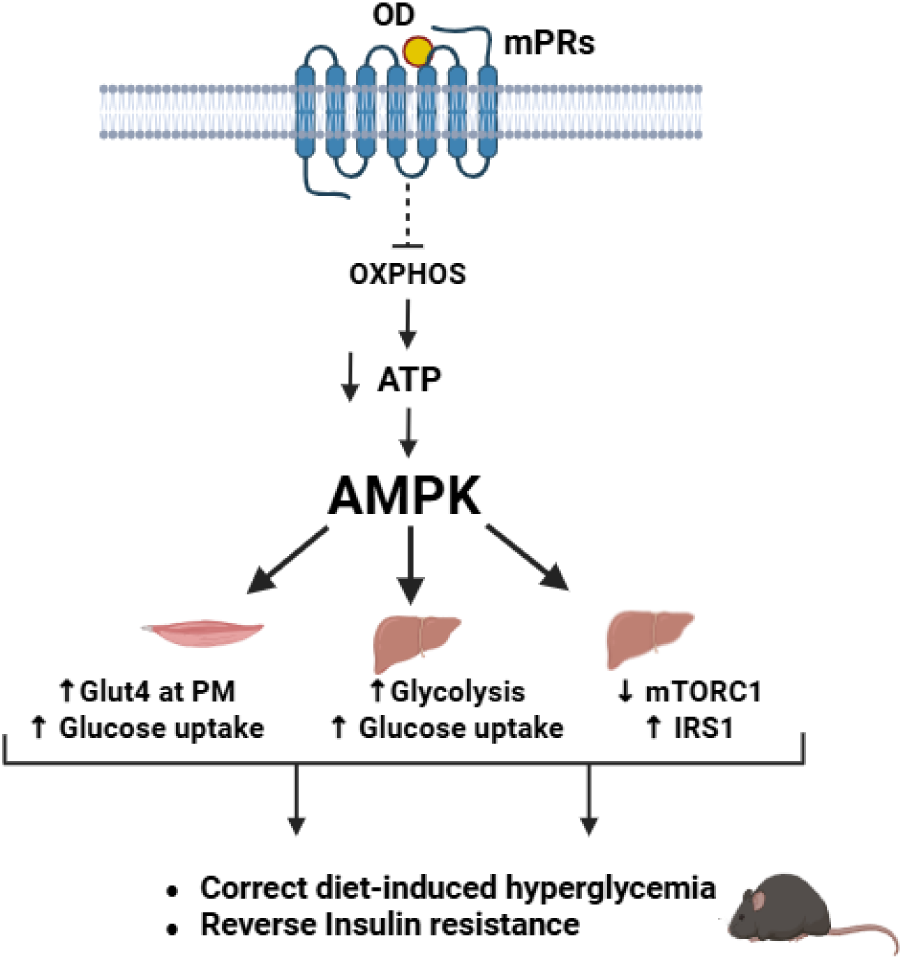
Proposed model for OD02-0 action in metabolic regulation. OD02-0 activates mPRs to promote AMPK signaling in both skeletal muscle and hepatocytes. In skeletal muscle, AMPK activation increases GLUT4 recruitment to the plasma membrane, enhancing glucose uptake. In hepatocytes, AMPK activation is associated with reduced oxidative phosphorylation and increased glycolytic flux, promoting glucose uptake, while suppression of mTORC1 is associated with preservation of IRS1 levels and improved insulin responsiveness. Together, these coordinated tissue-specific actions reverse diet-induced hyperglycemia and insulin resistance in vivo. Created with BioRender. Nader, N. (2025). https://BioRender.com/s0lexot

Previous studies have characterized OD02-0 as a selective mPR agonist with minimal activity at nuclear progesterone receptors (*27*). Since its initial characterization, OD02-0 has been widely used in diverse human and rodent cellular and tissue models, supporting its utility as a pharmacological tool to investigate mPR signaling (*35–40*). Consistent with this profile, we observed no evidence of transcriptional modulation in liver or muscle following treatment, supporting a predominantly non-genomic mode of action. In addition, a 28-day toxicological assessment in lean mice revealed no detectable adverse effects, further supporting its tolerability in preclinical models.

Interestingly, the signaling profile downstream of mPR activation partially parallels that of metformin, the current first-line therapy for T2DM. Both converge on AMPK activation and mTORC1 suppression, pathways known to enhance insulin sensitivity and regulate hepatic glucose metabolism (*65, 66*). Metformin requires OCT1-mediated uptake into hepatocytes, where its mitochondrial accumulation leads to Complex I inhibition and AMPK activation (*67, 68*), although such effects are most consistently observed at higher intracellular concentrations than those achieved under typical clinical conditions. In contrast, OD02-0 acts through membrane receptor-mediated signaling rather than intracellular accumulation and direct mitochondrial complex inhibition. This mechanistic distinction suggests that targeting mPR may provide a complementary strategy to existing therapies and may help address limitations associated with metformin’s gastrointestinal intolerance due to its high intestinal concentrations (*69*). In addition, progesterone signaling via mPR has been shown to influence incretin pathways, including GLP-1–mediated glucose control (*9*), raising the possibility that mPR activity intersects with established hormonal mechanisms relevant to diabetes treatment.

Several limitations warrant consideration. First, although OD02-0 demonstrates robust efficacy in diet-induced obese mice, its long-term durability and safety across additional models of diabetes remain to be determined. Second, while our findings establish convergence on AMPK-mTORC1 signaling, the precise upstream intermediates linking mPR activation to mitochondrial and glycolytic reprogramming require further investigation. Third, the relative contributions of individual mPR receptor subtypes were not dissected and may influence tissue-specific metabolic responses. More broadly, mPR family members exhibit context-dependent functions across neurological, vascular, reproductive, and oncologic settings (*10, 41–44, 70–74*). Studies in breast, prostate, pancreatic, and endometrial cancers suggest that mPR signaling can exert protective or pro-tumorigenic effects depending on cellular context, mPR subtypes and downstream pathway engagement. These observations underscore the importance of defining receptor subtype specificity, tissue-selective actions, and long-term systemic consequences before therapeutic application.

Advancing these findings toward clinical translation will require development of isoform-selective ligands, extended pharmacokinetic and toxicological profiling, validation in additional preclinical models, and assessment of combinatorial efficacy with established metabolic therapies.

Collectively, these results support membrane progesterone receptor signaling as a rapid endocrine-metabolic regulatory pathway that interfaces with central energy-sensing mechanisms to improve insulin sensitivity and glucose homeostasis. By expanding the framework of steroid hormone action beyond classical genomic pathways, this work identifies mPR as a promising molecular target in metabolic diseases characterized by insulin resistance and hyperglycemia.

## Acknowledgments

We are grateful to the genomic, proteomic and bioinformatics cores at Weill Cornell Medicine Qatar for their support in the transcriptomic and proteomic studies. We also thank Vivarium, histology and microscopy cores at WCMQ for their support in multiple experiments. The authors used ChatGPT (OpenAI, GPT-5.2) for grammar and language refinement. All scientific content, analysis, and interpretations were generated and verified by the authors. This work as well as the Cores are supported by the Biomedical Research Program at Weill Cornell Medical College in Qatar (BMRP) awarded to Khaled Machaca, a program funded by Qatar Foundation; and by NPRP-Standard (NPRP-S) thirteen (13th) Cycle grant 13S-0206-200274 from the Qatar National Research Fund (a member of Qatar Foundation), awarded to Nancy Nader. The statements made herein are solely the responsibility of the authors.

## Contribution statement

N. Nader conceived and designed the study.

N. Nader and K Machaca supervised the project and led data analysis and interpretation.

L. Zarif, S. Sherif, J. Al Hamaq, D. Al Qahtani, R. Courjaret, F. Yu, H. H Abunada and P. B. Vemulapalli performed experiments and acquired data.

S. Choi and F. Schmidt performed proteomic and phosphoproteomic analyses and contributed to data interpretation.

N. Nader drafted the manuscript. All authors contributed to data interpretation, critically revised the manuscript for important intellectual content, and approved the final version of the manuscript.

N. Nader and K. Machaca are the guarantors of this work and, as such, have full access to all the data in the study and take responsibility for the integrity of the data and the accuracy of the data analysis.

## Data and materials availability

The proteomic and phosphoproteomic datasets generated in this study have been deposited in the ProteomeXchange Consortium via the PRIDE partner repository under accession number PXD075864. During peer review, the datasets can be accessed using the following reviewer credentials: Username; Password: TrZdGIzJRziH. All other data supporting the findings of this study are available in the main text or the supplementary materials.

---

**Figure S1.**
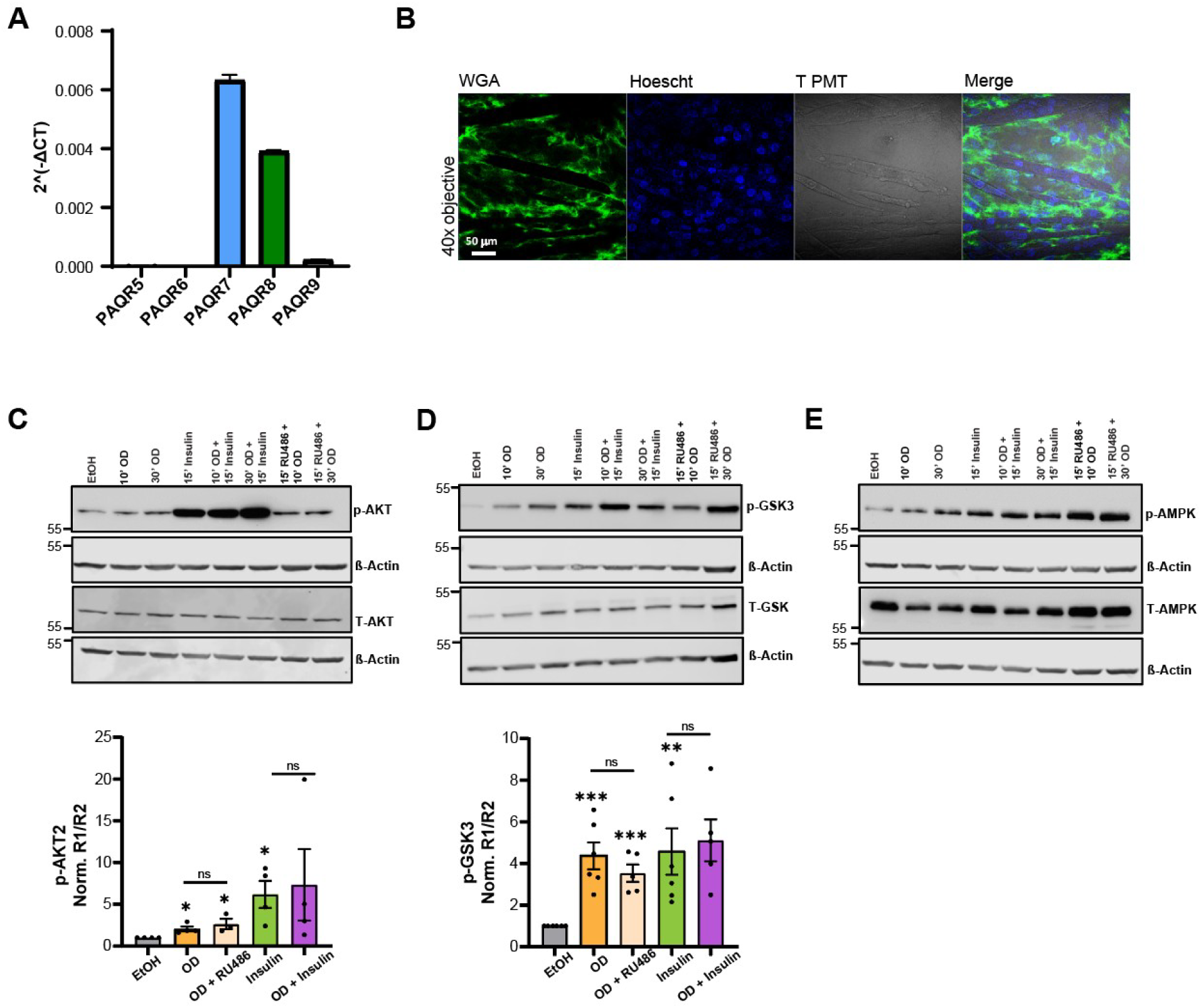
mPR signaling in C2C12 muscle cells. **(A)** mRNAs transcripts of PAQR5 for mPRγ, PAQR8 for mPRβ, PAQR7 for mPRα, PAQR6 for mPRδ and PAQR9 for mPRε using the Cycle threshold (Ct) generated from real time PCR, after correction on GAPDH levels as a loading control to obtain 2^(-ΔCT). **(B)** Fluorescence microscopy image of differentiated C2C12 myotubes showing their characteristic elongated morphology. Nuclei are stained with Hoechst (blue), and the plasma membrane is labeled with wheat germ agglutinin (WGA, green), confirming successful differentiation and membrane integrity. Scale bar is 50 μm. **(C/D/E)** *Upper panels*, representative Western blots for p-AKT/total AKT (C), p-GSK3β/total GSK3β (D), and pAMPK/total AMPK (E) in C2C12 myotubes treated with ethanol (EtOH), OD 02-0 (OD) for 10 or 30 min, insulin (15 min), OD + insulin, or OD with RU486 (a nuclear PR antagonist) for the time as indicated on the figure. β-Actin serves as the loading control. *Lower panels*, quantification of p-AKT/total AKT (C), and p-GSK3β/total GSK3β (D). Densitometric quantification was performed using ImageJ. For each sample, phospho/β-Actin (R1) and total/ β-Actin (R2) ratios were calculated, and phosphorylation was calculated as R1/R2. Final values were normalized to the vehicle (EtOH) control. Data represent mean ± SEM, n=3–5 biological replicates (unpaired two-tailed *t*-test. \**p* < 0.05; **p < 0.01; \*\*\**p* < 0.001; ns, not significant).

**Figure S2.**
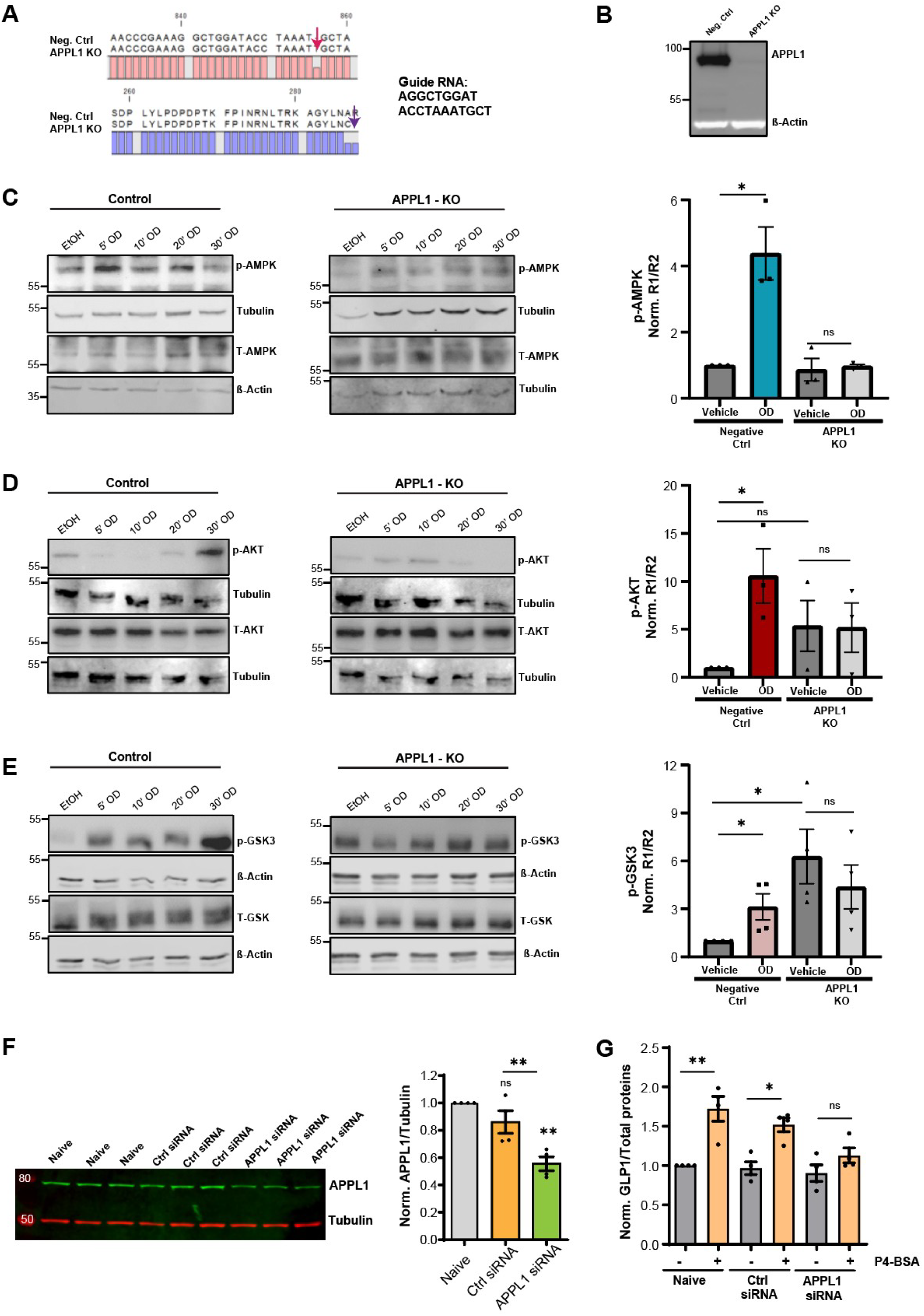
APPL1 is required for optimal mPR signaling and downstream metabolic responses. **(A)** C2C12 myotubes were transfected with a CRISPR/Cas9 construct targeting APPL1 using the guide RNA sequence AGGCTGGATACCTAAATGCT. Along with the guide RNA sequence, panel A shows the aligned cDNA (*top panel*) and translated protein sequences (*bottom panel*) comparing negative control (Neg. Ctrl) and APPL1 knockout (APPL1 KO) cells. In the KO cells, a single-nucleotide insertion at the guide RNA target site (indicated by the pink arrow) results in a frameshift, leading to the introduction of a premature stop codon (purple arrow) and truncation of the APPL1 protein. **(B)** Representative Western blot confirming the loss of APPL1 protein in C2C12 cells from the knockout clones, compared to control cells. β-Actin was used as a loading control. **(C/D/E)** *Left panels*, representative Western blot for p-AMPK/total AMPK (C), p-AKT/total AKT (D) and pGSK3β/total GSK3β (E) in negative control (Control) and APPL1 knockout (APPL1-KO) C2C12 myotubes treated with ethanol (EtOH, vehicle), OD 02-0 (OD) for 5-, 10-, 20- and 30-min. Tubulin or β-Actin served as loading controls. *Right panels*, densitometric quantification for p-AMPK (C), p-AKT (D) and p-GSK3β (E) was performed using ImageJ. For each sample, phospho/loading control (R1) and total/loading control (R2) ratios were calculated, and phosphorylation was calculated as R1/R2. Final values were normalized to the vehicle (EtOH) from negative control cells. Although multiple time points were tested for OD (5, 10, 20, and 30 minutes), for each experiment, only the time point showing the maximum effect on phosphorylation of AKT2, GSK3β and AMPK was included in the quantification graph. This approach accounts for temporal variability in response across replicates. Data represent mean ± SEM from n=3 to 4 independent experiments (unpaired two-tailed *t*-test. \**p* < 0.05; **p < 0.01; ns, not significant. **(F)** Representative Western blot (*left panel*) and densitometric quantification (*right panel*) showing effective APPL1 knockdown using siRNA in Glutag cells, compared to naïve and control siRNA-treated conditions. Data represent mean ± SEM from *n*=4 independent experiments (unpaired two-tailed *t*-test. **p < 0.01; ns, not significant). **(G)** Quantification of GLP-1 protein levels in naïve, control siRNA, and APPL1 siRNA-treated Glutag cells, with or without membrane-impermeable P4-BSA stimulation. GLP-1 levels were normalized to total protein content (cells + media) and expressed relative to the naïve condition. Data represent mean ± SEM from *n*=4 independent experiments (one-way ANOVA. *p < 0.05; **p < 0.01; ns, not significant).

**Figure S3.**
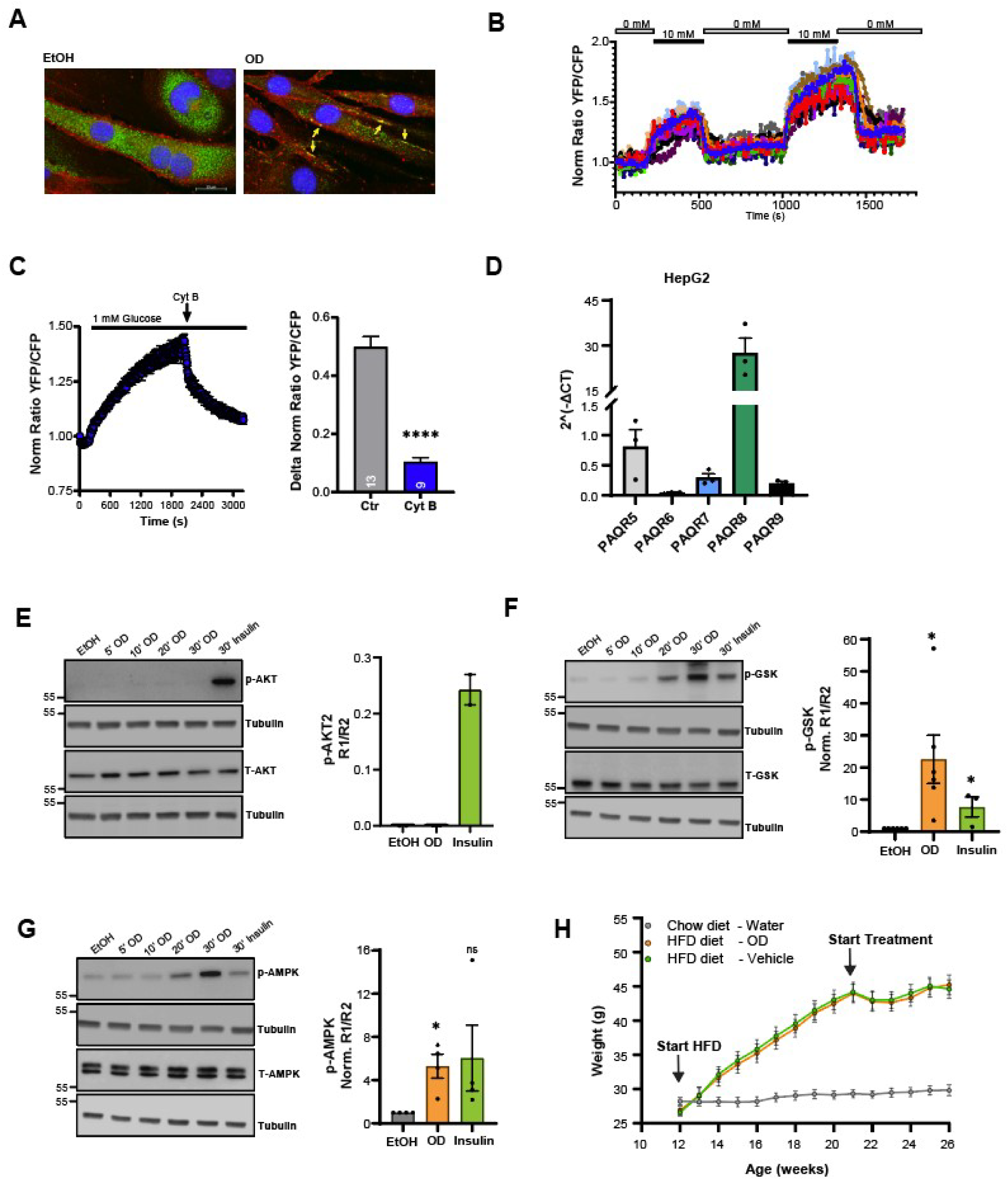
Glucose uptake in C2C12 and OD 02-0 signaling in HepG2 cells. **(A)** Representative confocal microscopy images showing subcellular localization of endogenous GLUT4 (green) and Na⁺/K⁺ pump (red) in C2C12 myotubes treated with vehicle (EtOH) or OD 02-0 (OD). Nuclei were counterstained with Hoechst (blue). Merged images highlight GLUT4 translocation to the membrane following OD treatment. Scale bar is 50 μm. **(B)** YFP/CFP fluorescence ratio traces per individual cell over time showing FRET-based glucose sensor responses under basal (0 mM) and glucose-stimulated (10 mM) conditions. **(C)** *Left panel:* FRET traces in C2C12 myotubes exposed to 1 mM glucose before and after addition of cytochalasin B (Cyt B) that block GLUT4 translocation. *Right panel*: quantification of glucose uptake based on FRET ratio changes (ΔFRET) between untreated or after cytochalasin addition. Data represent mean ± SEM; 9 to 13 cells per group (unpaired two-tailed *t*-test. \*\*\**p* < 0.001). **(D)** mRNAs transcripts of PAQR5 for mPRγ, PAQR8 for mPRβ, PAQR7 for mPRα, PAQR6 for mPRδ and PAQR9 for mPRε using the Cycle threshold (Ct) generated from real time PCR, after correction on GAPDH levels as a loading control to obtain 2^(-ΔCT). **(E/F/G)** OD 02-0 induces phosphorylation of key metabolic signaling proteins in human HepG2 cells. *Left panels:* representative Western blots for phospho- and total AKT (E), GSK3β (F), and AMPK (G) following treatment with ethanol (EtOH, for 30 min), OD 02-0 (OD, for 5, 10, 20 and 30 min), or insulin (for 30 min). Tubulin was used as loading control. *Right panels:* densitometric quantification performed using ImageJ. For each sample, phospho/tubulin (R1) and total/tubulin (R2) ratios were calculated, and phosphorylation was calculated as R1/R2. For GSK3β and AMPK, final values were normalized to the vehicle (EtOH) control. For phospho-AKT2, only raw ratios are presented, as no detectable signal was observed in the EtOH-treated condition for normalization. Although multiple time points were tested for OD (5, 10, 20, and 30 minutes), for each of the three independent experiments, only the time point showing the maximum effect on phosphorylation of AKT2, GSK3β and AMPK was included in the quantification graph. This approach accounts for temporal variability in response across replicates. Data represent mean ± SEM from n = 3 independent experiments (unpaired two-tailed *t*-test. \**p* < 0.05; **p < 0.01; ns, not significant). **(H)** Body weight progression during high-fat diet and OD02-0 treatment. Male mice were placed on a high-fat diet (HFD) starting at 12 weeks of age (arrow). After ∼8 weeks on HFD, mice were randomized to receive OD02-0 or vehicle, while chow-fed controls received water (arrow: start of treatment). Body weight was measured weekly. Both HFD groups gained comparable weight throughout the study, with no significant differences between OD02-0- and vehicle-treated mice. Data represent mean ± SEM.

**Figure S4.**
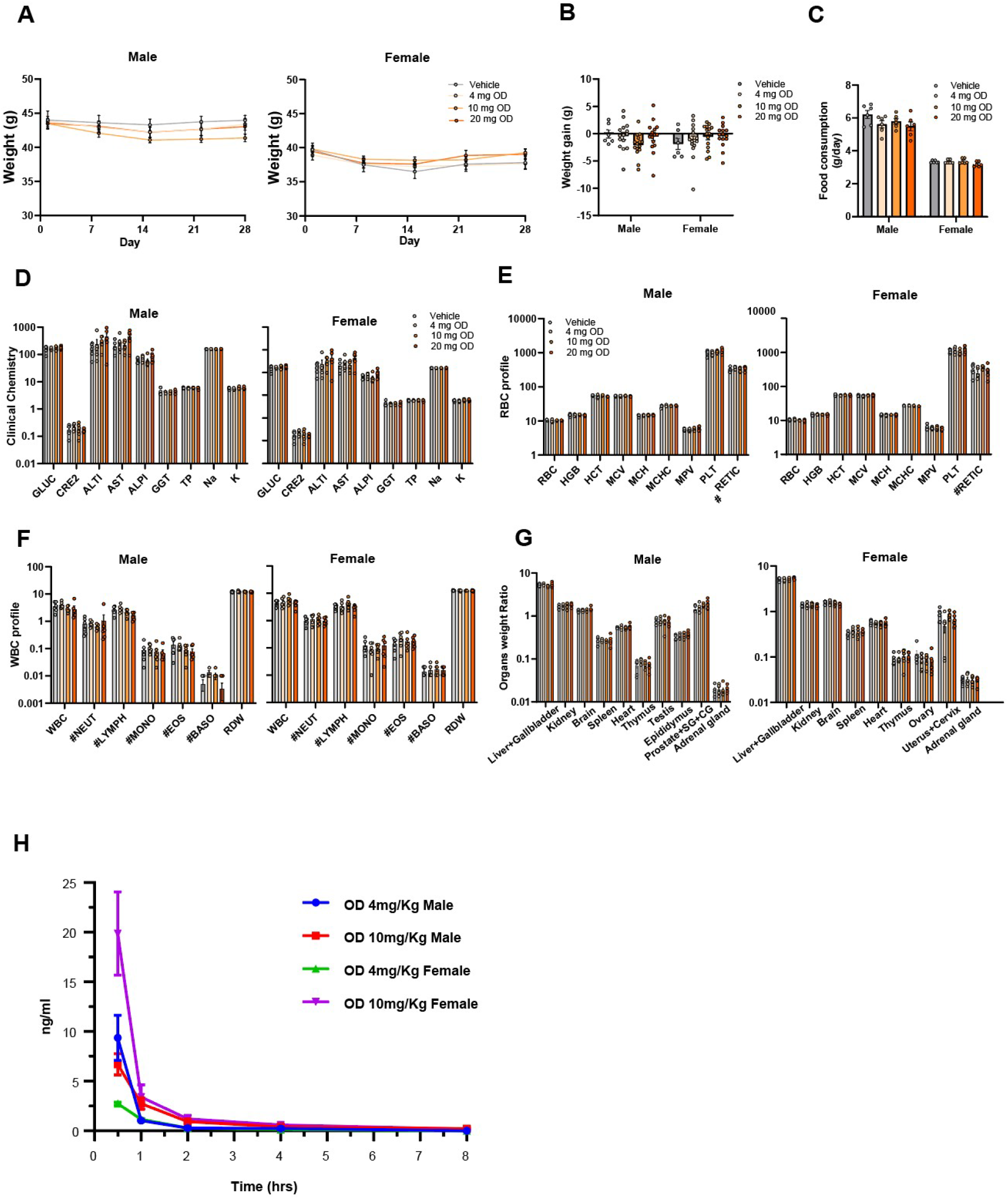
*In vivo* safety profile of chronic OD administration in male and female mice. **(A)** Body weight measurements over time in male (*left panel*) and female (*right panel*) mice across treatment groups, monitored during the 28-day OD treatment. Data are presented as mean ± SEM; n=6 mice per group. **(B)** Average body weight gain in male and female mice across treatment groups at endpoint. Data are presented as mean ± SEM; n=6 mice per group. **(C)** Food consumption in male and female mice across treatment groups at endpoint. Data are presented as mean ± SEM; n=6 mice per group. **(D)** Serum chemistry analysis for male (*left panel*) and female (*right panel*) mice, including glucose (GLUC), creatinine (CRE2), liver enzymes (ALT, AST, ALP, GGT), total protein (TP), and electrolytes (Na, K), showing no major toxicity signals across treatment groups. Data are presented as mean ± SEM; n=6 mice per group (unpaired twotailed *t*-test). **(E/F)** Red (RBC, E) and white (WBC, F) blood cells indices in male (*left panels*) and female (*right panels*) mice after 4-week daily treatment with vehicle or OD at 4, 10, or 20 mg/kg. Parameters measured include RBC count, hemoglobin (HGB), hematocrit (HCT), mean corpuscular volume (MCV), mean corpuscular hemoglobin (MCH), MCH concentration (MCHC), mean platelet volume (MPV), platelet count (PLT), and reticulocyte number (#RETIC), neutrophils (#NEUT), lymphocytes (#LYMPH), monocytes (#MONO), eosinophils (#EOS), basophils (#BASO), and red cell distribution width (RDW). Data are presented as mean ± SEM; n=6 mice per group (unpaired two-tailed *t*test). **(G)** Relative organ weight ratios (organ weight/body weight) for major organs in male (*left panel*) and female (*right panel*) mice, including liver and gallbladder, kidneys, brain, spleen, heart, thymus, testis, epididymis, prostate/seminal/gland complex, adrenal glands, ovaries, and uterus/cervix. Data are presented as mean ± SEM; n=6 mice per group. Statistical analysis: unpaired two-tailed *t*-test. **(H)** Plasma pharmacokinetics of OD 02-0 in male and female mice. Plasma concentrations of OD 02-0 were measured over an 8-hour period following a single intraperitoneal injection of either 4 mg/kg or 10 mg/kg in male and female mice. OD 02-0 exhibited rapid absorption, with peak plasma levels occurring within the first hour, followed by a sharp decline and near-baseline levels by 4–6 hours across all groups. Female mice showed higher initial plasma exposure at the 10 mg/kg dose compared with males, whereas elimination profiles were comparable. Data are presented as mean ± SEM.

**Figure S5.**
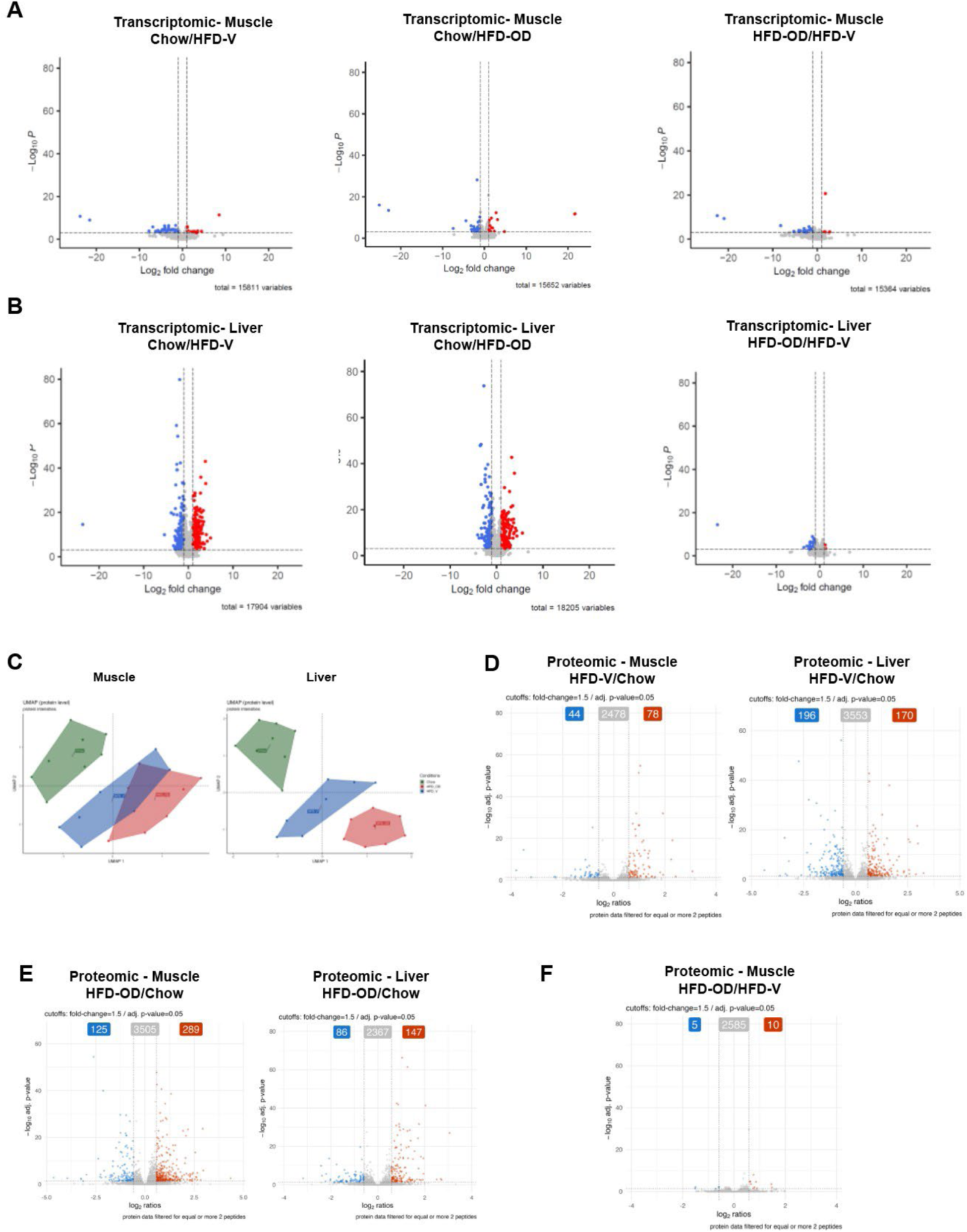
Multi-omics profiling from muscle and liver tissues collected from mice. **(A/B)** Volcano plots of differentially expressed genes in muscle (A) and liver (B) comparing Chow vs. HFD-V (*left*), Chow vs. HFD-OD (*middle*), and HFD-OD vs HFD-V (*right*). Significantly upregulated genes are shown in red and downregulated genes in blue (cutoff: fold change >1.5 or <−1.5; adjusted p < 0.05). **(C)** Uniform Manifold Approximation and Projection (UMAP) analysis of the transcriptomic data from muscle (*left*) and liver (*right*). Each point represents a sample, with clustering observed by condition (Chow, HFD-V, and HFDOD), indicating distinct transcriptional profiles. **(D/E/F)** Volcano plots of differentially expressed proteins in muscle (*left*) or liver (*right*) comparing HFD-V vs. Chow (D) and HFD-OD vs. Chow (E) in both tissues, and HFD-OD vs. Chow in muscle (F) . Significantly upregulated proteins are shown in red and downregulated in blue (cutoff: fold change >1.5 or <-1.5; p < 0.05). The number of significant proteins is indicated.

**Figure S6.**
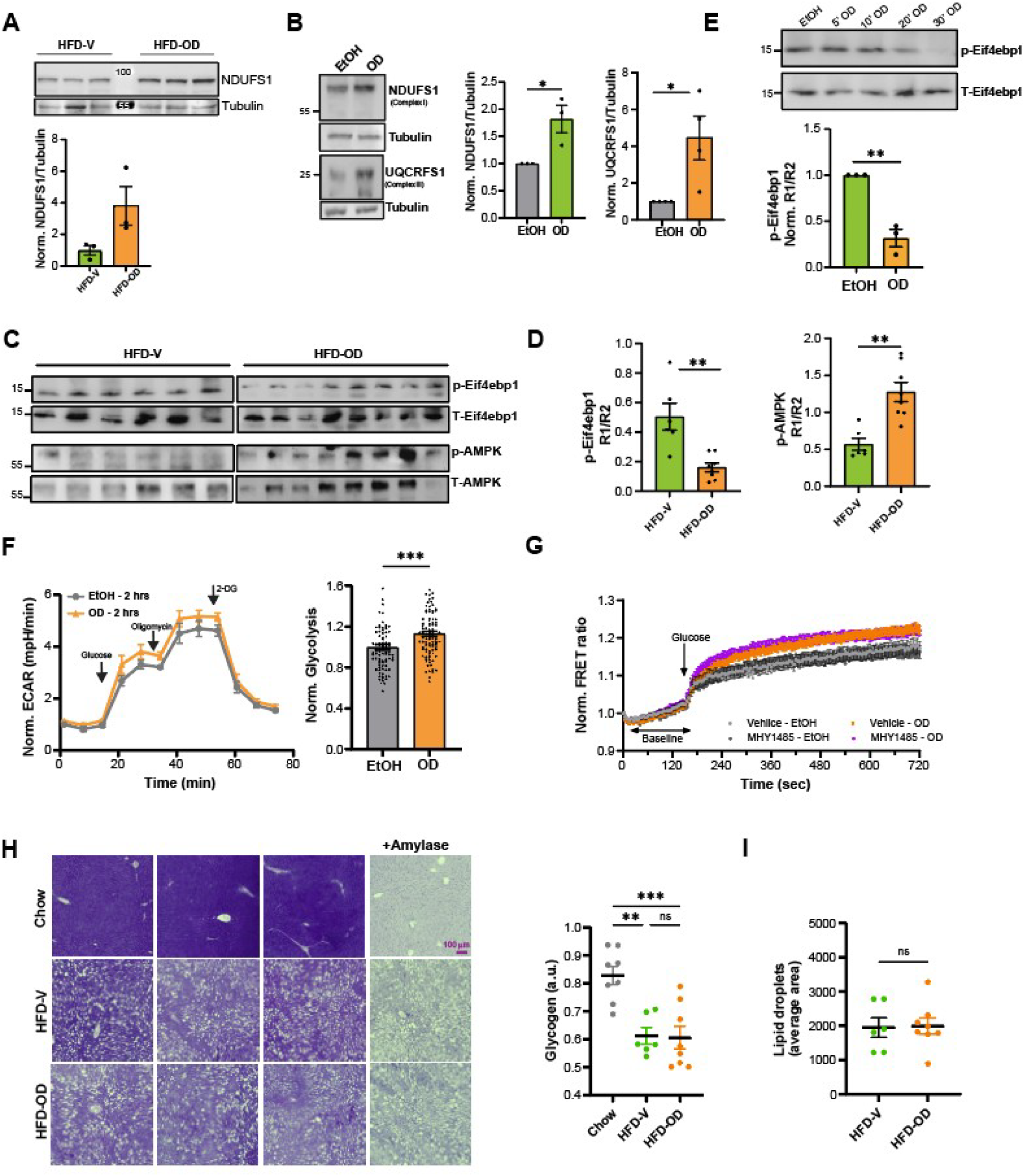
Exploration of metabolic pathway changes. **(A)** Representative Western blot (*upper*) and quantification (*lower)* of NDUFS1 (Complex I subunit) protein levels in liver tissues from HFD-fed mice treated with vehicle (HFD-V) or OD02-0 (HFD-OD). Tubulin served as a loading control. For each sample, the ratio of NDUFS1 over tubulin was calculated and normalized to the average ratios of the HFD-V samples. Data are presented as mean ± SEM; n=3 mice per group. **(B)** Representative Western blot (*left*) and quantification (*right*) of NDUFS1 (Complex I subunit) and UQCRFS1 (Complex III) in HepG2 cells treated with ethanol (EtOH) or OD 02-0 (OD) for 3 days. Tubulin was used as a loading control. For each sample, the ratio of NDUFS1 and UQCRFS1 over tubulin were calculated and normalized to the average of the ratios of the EtOH condition. Data are presented as mean ± SEM; n=3 to 4 independent experiments (unpaired two-tailed *t*-test. *p < 0.05). **(C)** Representative Western blots for liver proteins extracts from HFD-fed mice showing increased phosphorylation of AMPK (Thr172) and decreased phosphorylation of Eif4ebp1 in OD 02-0-treated mice compared to vehicle. Tubulin or β-actin was used as loading control (not shown). **(D)** Densitometric quantification of Western blots shown in (C). For each sample, phospho/loading control and total/loading control ratios were calculated (R1 and R2, respectively), and phosphorylation was expressed as R1/R2. Data represent mean ± SEM; n=6 to 8 mice per group, and WB was repeated twice. Statistical analysis: unpaired twotailed *t*-test. *\*\**p < 0.01. **(E)** *Upper*, representative Western blot showing Eif4ebp1 phosphorylation levels in HepG2 cells treated with EtOH (vehicle) or OD 02-0 for 5, 10, 20, or 30 minutes. β-Actin was used as loading control. *Lower*, densitometric quantification was performed using ImageJ. For each sample, phosphor-Eifebp4/β-actin (R1) and total-Eifebp4/βactin (R2) were calculated, and phosphorylation was expressed as R1/R2. Final values were normalized to the vehicle (EtOH) control. Although multiple time points were tested (5, 10, 20, and 30 minutes), for each of the three independent experiments, only the time point showing the maximum inhibition of Eif4ebp1 phosphorylation by OD 02-0 was included in the quantification graph. This approach accounts for temporal variability in response across replicates. Data represent mean ± SEM from n=3 independent experiments (unpaired two-tailed *t*-test. **p < 0.01). **(F)** Glycolytic function was assessed in HepG2 cells treated with OD02-0 (10⁻⁶ M) or vehicle (EtOH) for 2 hours, using the Seahorse XF Glycolysis Stress Test. *Left,* extracellular acidification rate (ECAR) was measured over time in response to sequential injections of glucose, oligomycin, and 2-deoxyglucose. *Right*, normalized glycolytic rate in HepG2 cells treated with OD 02-0 (10⁻⁶ M) or vehicle (EtOH, 0.1%). Data represent mean ± SEM from 3 independent experiments (unpaired two-tailed *t*-test. **p < 0.01). **(G)** Time course of normalized FRET ratio following glucose addition in HepG2 cells expressing the FLII¹²Pglu-700 μδ6 glucose biosensor. Cells were pretreated with MHY1485 (10 μM) or vehicle (DMSO 0.1%) for 30 min, followed by stimulation with OD 02-0 (10⁻⁶ M) or vehicle (EtOH 0.1%) for 30 min. **(H)** *Left*, representative periodic acid–Schiff (PAS) staining images of liver sections from chow, HFD-V, and HFDOD treated mice. Amylase was used as a glycogen-specific control. Scale bar, 100 μm. *Right*, quantification of hepatic glycogen levels in chow, HFD-V, and HFD-OD mice. Data are presented as mean ± SEM; n=6 to 8 mice per group (one-way Anova test. \*\**p* < 0.01 \*\*\**p* < 0.001; ns, not significant). **(I)** Quantification of hepatic lipid droplet average area in HFD-V and HFD-OD mice. Data represent mean ± SEM; n=6 to 8 mice per group (unpaired two-tailed *t*-test. ns, not significant).

**Table S1.** Toxicokinetic Parameters of Org OD02-0 Following Oral Gavage of Org OD02-0 in CD1 Mice on Day 1 and Day 28.

| Analyte | Sex | Day Nominal | Treatment | Dose<br>(mg/kg/day) | Cmax<br>(ng/m) | Tmax<br>(h) | T1/2<br>(h) |
| --- | --- | --- | --- | --- | --- | --- | --- |
| Org OD02-0 | Male | Day 1 | Org OD02-0 | 4 | 49.29 | 1.0 | NC |
|  |  |  |  | 10 | 70.87 | 0.5 | 4.7 |
|  |  |  |  | 20 | 68.77 | 0.5 | NC |
| Org OD02-0 | Female | Day 1 | Org OD02-0 | 4 | 46.30 | 1.0 | 3.4 |
|  |  |  |  | 10 | 37.07 | 0.5 | 8.1 |
|  |  |  |  | 20 | 79.11 | 0.5 | 3.8 |
| Org OD02-0 | Sex<br>Combined | Day 1 | Org OD02-0 | 4 | 47.80 | 1.0 | NC |
|  |  |  |  | 10 | 53.97 | 0.5 | 6.4 |
|  |  |  |  | 20 | 73.94 | 0.5 | NC |
| Org OD02-0 | Male | Day 28 | Org OD02-0 | 4 | 1.91 | 0.5 | 2.3 |
|  |  |  |  | 10 | 11.89 | 1.0 | NC |
|  |  |  |  | 20 | 46.30 | 0.5 | NC |
| Org OD02-0 | Female | Day 28 | Org OD02-0 | 4 | 3.53 | 0.5 | NC |
|  |  |  |  | 10 | 41.31 | 1.0 | 1.6 |
|  |  |  |  | 20 | 44.12 | 0.5 | 1.2 |
| Org OD02-0 | Sex<br>Combined | Day 28 | Org OD02-0 | 4 | 2.72 | 0.5 | NC |
|  |  |  |  | 10 | 26.60 | 1.0 | NC |
|  |  |  |  | 20 | 45.21 | 0.5 | NC |

**Table S2.** List of organs assessed by histopathology.

|  | **Tissue** |  | **Tissue** |
| --- | --- | --- | --- |
| 1. | Artery, Aorta | 27. | Large Intestine, Rectum |
| 2. | Bone, Femur/Joint, Femorotibial | 28. | Liver |
| 3. | Bone, Sternum | 29. | Lung |
| 4. | Brain | 30. | Lymph node, Mesenteric |
| 5. | Cervix | 31. | Lymph node, Mandibular |
| 6. | Epididymis | 32. | Muscle, Skeletal |
| 7. | Esophagus | 33. | Nerve, Optic |
| 8. | Eye | 34. | Nerve, Sciatic |
| 9. | Gall bladder | 35. | Ovary |
| 10. | Gland, Adrenal | 36. | Oviduct |
| 11. | Gland, Coagulating | 37. | Pancreas |
| 12. | Gland, Harderian | 38. | Skin |
| 13. | Gland, Mammary | 39. | Small intestine, Duodenum |
| 14. | Gland, Parathyroid | 40. | Small intestine, Ileum |
| 15. | Gland, Pituitary | 41. | Small intestine, Jejunum |
| 16. | Gland, Prostate | 42. | Spinal cord |
| 17. | Gland, Salivary (mandibular, parotid and sublingual) | 43. | Spleen |
| 18. | Gland, Seminal vesicle | 44. | Stomach |
| 19. | Gland, Thyroid | 45. | Testis |
| 20. | Gross lesion | 46. | Thymus |
| 21. | Head | 47. | Tongue |
| 22. | Heart | 48. | Trachea |
| 23. | Kidney | 49. | Urinary bladder |
| 24. | Large intestine, Cecum | 50. | Vagina |
| 25. | Large intestine, Colon | 51. | Uterus |

**Table S3.**
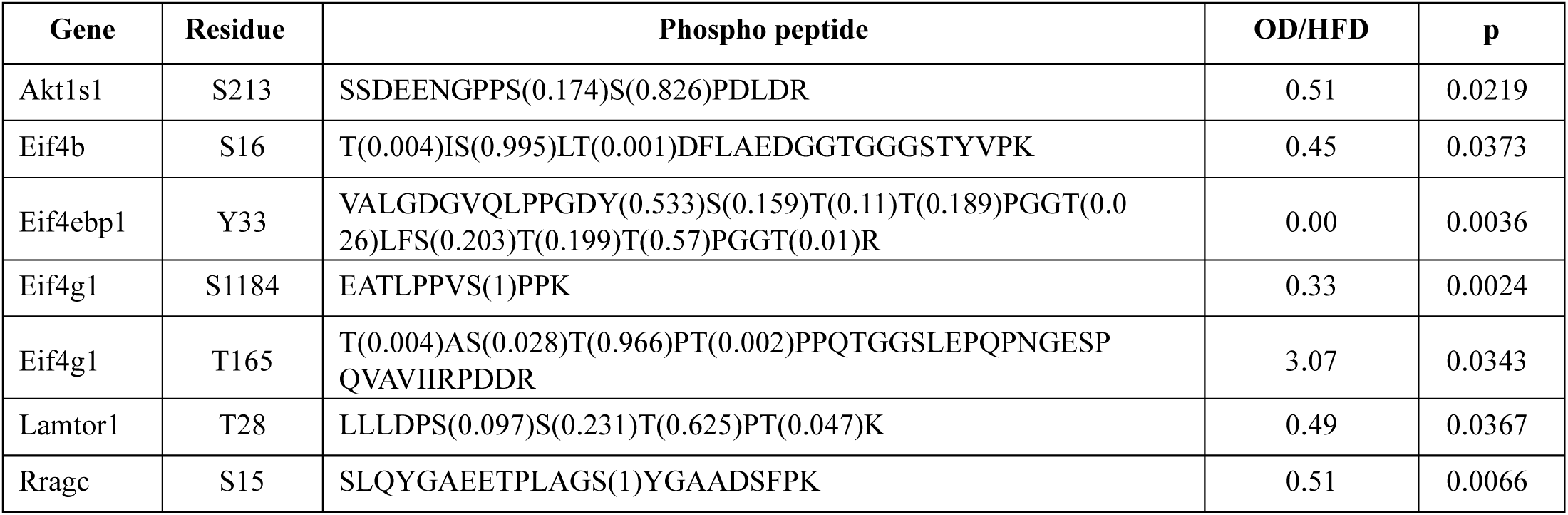
Phosphopeptides - mTOR pathway.

**Table S4.** Phosphopeptides - glycogen metabolism pathway.

| Gene | Residue | Phospho peptide | OD/HFD | p |
| --- | --- | --- | --- | --- |
| Pgm1 | T357 | IALYETPT(1)GWK | 0.45 | 0.0017 |
| Gys2 | S627 | FHLEPTS(1)PPTTDGFK | 0.26 | 0.0442 |
| Phkb | S19 | S(0.066)GS(0.933)VY(0.001)EPLK | 0.51 | 0.0405 |
| Calm3 | S102 | DGNGYIS(1)AAELR | 0.30 | 0.0266 |
| Ppp2r5d | S44 | DGGGENTDEAQPQPQS(0.107)QS(0.678)PS(0.107)S(0.107)NK | 0.32 | 0.0354 |

**Supp. Table S5.**
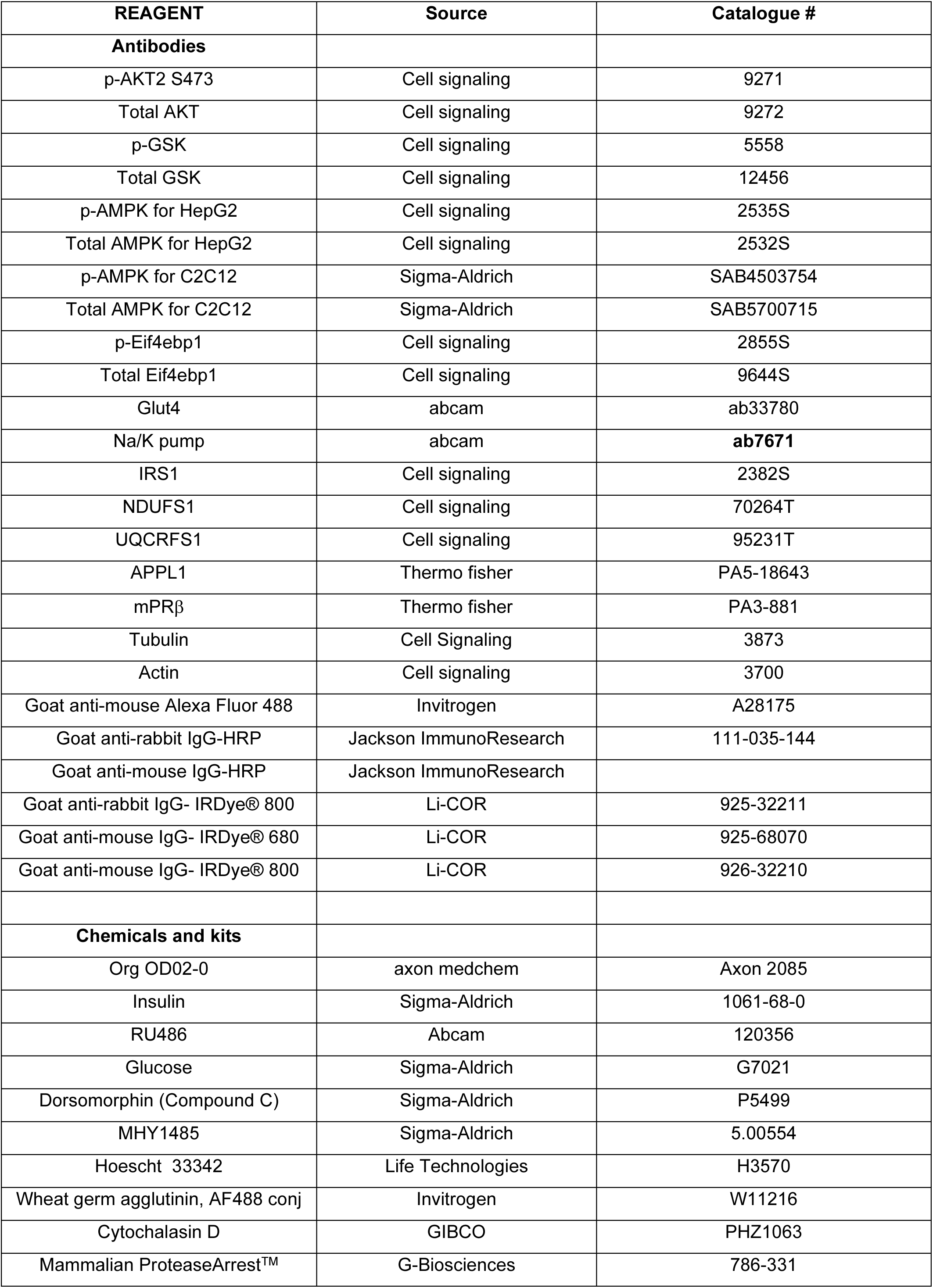
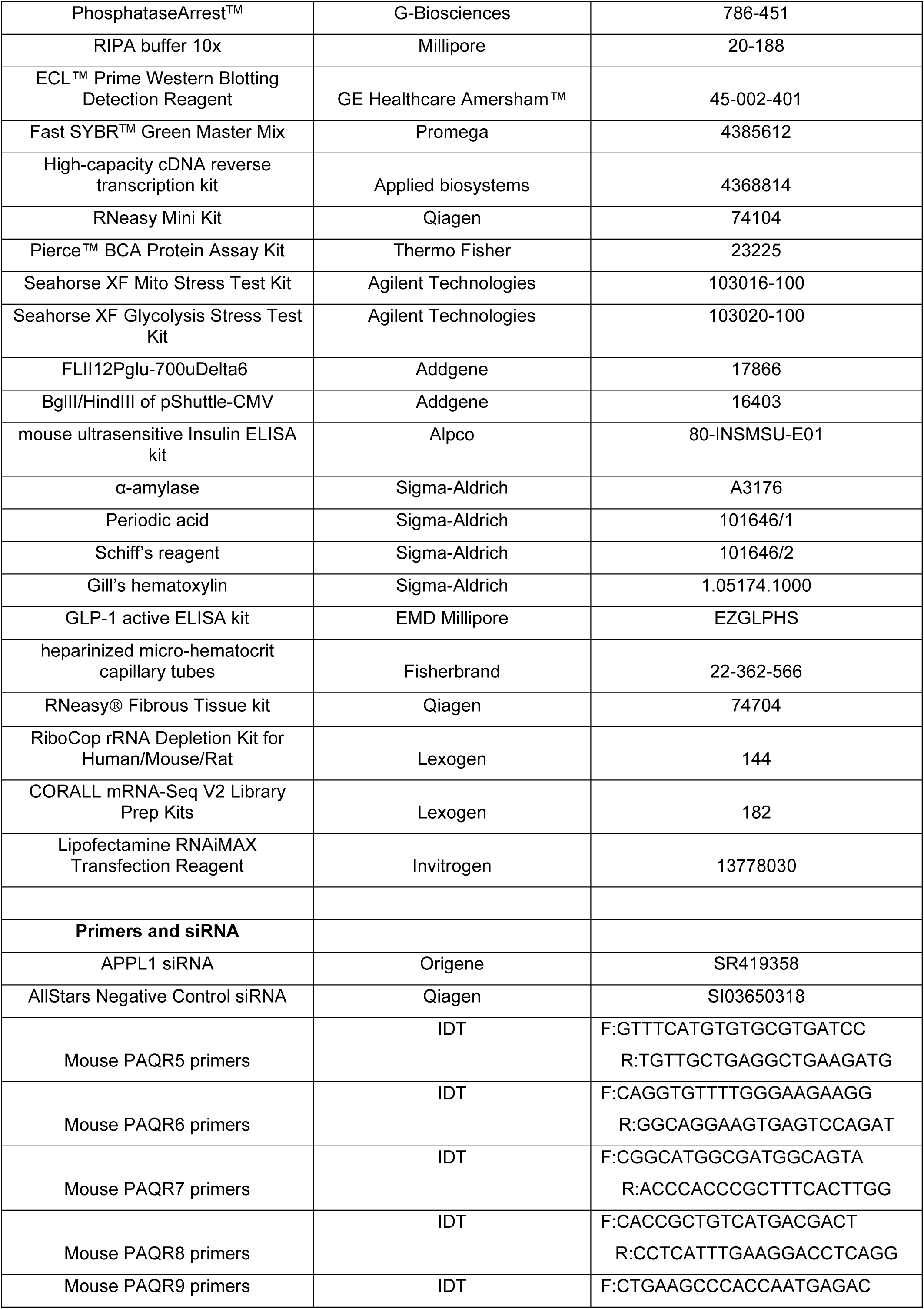
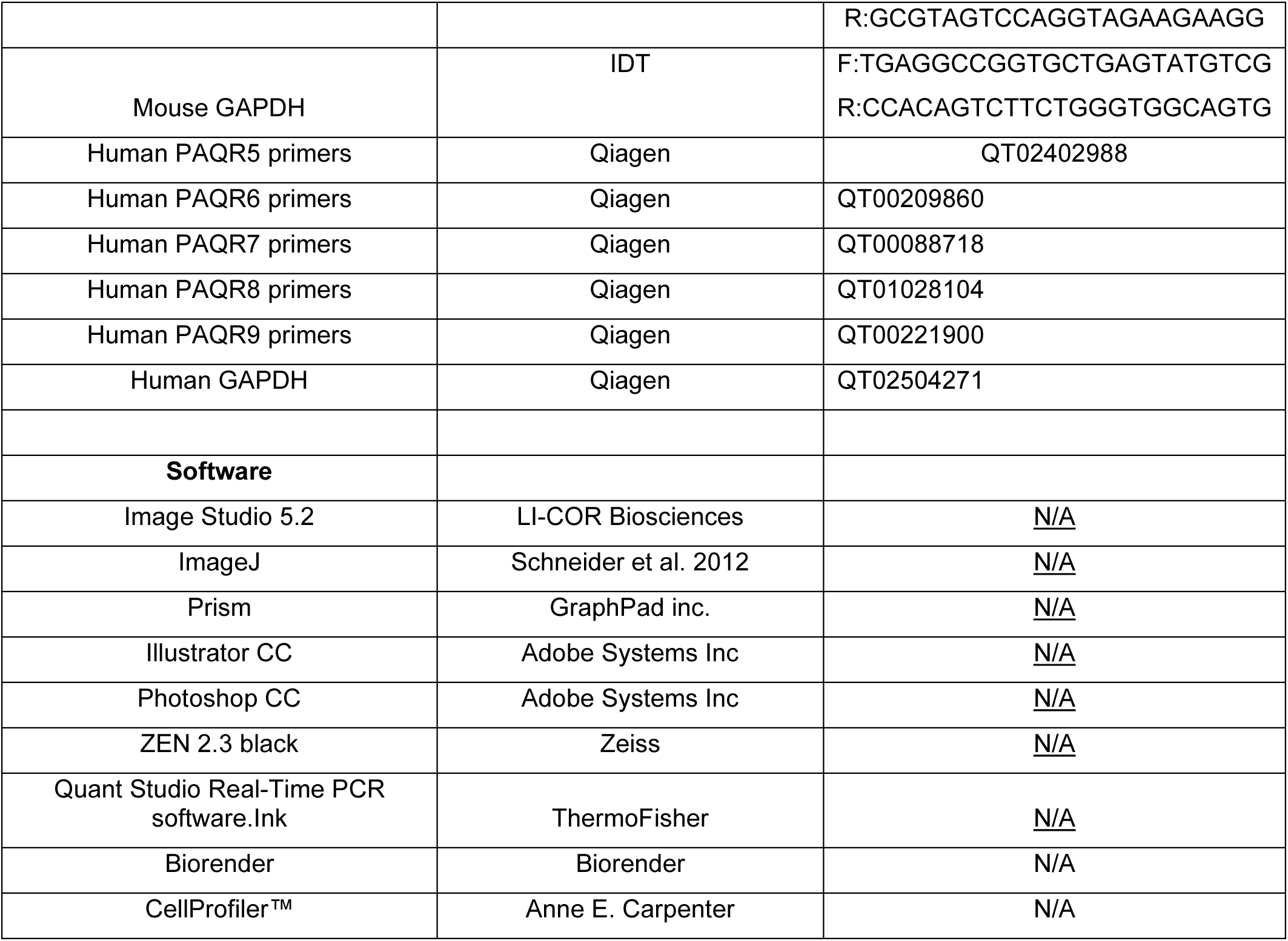
List of antibodies, chemicals, reagents, and oligoes.

## References

1. D. Garg, S. S. M. Ng, K. M. Baig, P. Driggers, J. Segars, Progesterone-Mediated Non-Classical Signaling. Trends Endocrinol Metab 28, 656–668 (2017).

2. P. Thomas, Rapid steroid hormone actions initiated at the cell surface and the receptors that mediate them with an emphasis on recent progress in fish models. Gen Comp Endocrinol 175, 367–383 (2012).

3. G. E. Dressing, J. E. Goldberg, N. J. Charles, K. L. Schwertfeger, C. A. Lange, Membrane progesterone receptor expression in mammalian tissues: a review of regulation and physiological implications. Steroids 76, 11–17 (2011).

4. R. Rekawiecki, M. K. Kowalik, J. Kotwica, Nuclear progesterone receptor isoforms and their functions in the female reproductive tract. Pol J Vet Sci 14, 149–158 (2011).

5. Y. T. Tang et al., PAQR proteins: a novel membrane receptor family defined by an ancient 7-transmembrane pass motif. J Mol Evol 61, 372–380 (2005).

6. P. Moussatche, T. J. Lyons, Non-genomic progesterone signalling and its non-canonical receptor. Biochem Soc Trans 40, 200–204 (2012).

7. P. Valadez-Cosmes, E. R. Vazquez-Martinez, M. Cerbon, I. Camacho-Arroyo, Membrane progesterone receptors in reproduction and cancer. Mol Cell Endocrinol 434, 166–175 (2016).

8. C. Dosiou et al., Expression of membrane progesterone receptors on human T lymphocytes and Jurkat cells and activation of G-proteins by progesterone. J Endocrinol 196, 67–77 (2008).

9. G. B. Flock, X. Cao, M. Maziarz, D. J. Drucker, Activation of enteroendocrine membrane progesterone receptors promotes incretin secretion and improves glucose tolerance in mice. Diabetes 62, 283–290 (2013).

10. A. Wendler, M. Wehling, Many or too many progesterone membrane receptors? Clinical implications. Trends Endocrinol Metab 33, 850–868 (2022).

11. P. Thomas et al., Steroid and G protein binding characteristics of the seatrout and human progestin membrane receptor alpha subtypes and their evolutionary origins. Endocrinology 148, 705–718 (2007).

12. N. Nader et al., Membrane progesterone receptor induces meiosis in Xenopus oocytes through endocytosis into signaling endosomes and interaction with APPL1 and Akt2. PLoS Biol 18, e3000901 (2020).

13. N. Nader et al., Progesterone induces meiosis through two obligate co-receptors with PLA2 activity. Elife 13, (2025).

14. N. Nader, R. Courjaret, M. Dib, R. P. Kulkarni, K. Machaca, Release from Xenopus oocyte prophase I meiotic arrest is independent of a decrease in cAMP levels or PKA activity. Development 143, 1926–1936 (2016).

15. E. Maury, S. M. Brichard, Adipokine dysregulation, adipose tissue inflammation and metabolic syndrome. Mol Cell Endocrinol 314, 1–16 (2010).

16. E. Nigro et al., New insight into adiponectin role in obesity and obesity-related diseases. Biomed Res Int 2014, 658913 (2014).

17. R. W. Mackenzie, B. T. Elliott, Akt/PKB activation and insulin signaling: a novel insulin signaling pathway in the treatment of type 2 diabetes. Diabetes Metab Syndr Obes 7, 55–64 (2014).

18. K. K. Cheng et al., APPL1 potentiates insulin-mediated inhibition of hepatic glucose production and alleviates diabetes via Akt activation in mice. Cell Metab 9, 417–427 (2009).

19. K. K. Cheng, K. S. Lam, B. Wang, A. Xu, Signaling mechanisms underlying the insulin-sensitizing effects of adiponectin. Best Pract Res Clin Endocrinol Metab 28, 3–13 (2014).

20. L. Zhou et al., Adiponectin activates AMP-activated protein kinase in muscle cells via APPL1/LKB1-dependent and phospholipase C/Ca2+/Ca2+/calmodulin-dependent protein kinase kinase-dependent pathways. J Biol Chem 284, 22426–22435 (2009).

21. G. Zhou et al., Role of AMP-activated protein kinase in mechanism of metformin action. The Journal of clinical investigation 108, 1167–1174 (2001).

22. D. J. Drucker, Efficacy and Safety of GLP-1 Medicines for Type 2 Diabetes and Obesity. Diabetes Care, (2024).

23. H. Takanaga, B. Chaudhuri, W. B. Frommer, GLUT1 and GLUT9 as major contributors to glucose influx in HepG2 cells identified by a high sensitivity intramolecular FRET glucose sensor. Biochim. Biophys. Acta 1778, 1091–1099 (2008).

24. T. C. He et al., A simplified system for generating recombinant adenoviruses. Proc Natl Acad Sci U S A 95, 2509–2514 (1998).

25. D. R. Stirling et al., CellProfiler 4: improvements in speed, utility and usability. BMC Bioinformatics 22, 433 (2021).

26. S. Michalik et al., SpectroPipeR-a streamlining post Spectronaut® DIA-MS data analysis R package. Bioinformatics 41, (2025).

27. J. Kelder et al., Comparison between steroid binding to membrane progesterone receptor alpha (mPRalpha) and to nuclear progesterone receptor: correlation with physicochemical properties assessed by comparative molecular field analysis and identification of mPRalpha-specific agonists. Steroids 75, 314–322 (2010).

28. R. T. Watson, J. E. Pessin, Intracellular organization of insulin signaling and GLUT4 translocation. Recent Prog Horm Res 56, 175–193 (2001).

29. J. R. Jaldin-Fincati, M. Pavarotti, S. Frendo-Cumbo, P. J. Bilan, A. Klip, Update on GLUT4 Vesicle Traffic: A Cornerstone of Insulin Action. Trends Endocrinol Metab 28, 597–611 (2017).

30. M. E. Meyer et al., Agonistic and antagonistic activities of RU486 on the functions of the human progesterone receptor. Embo j 9, 3923–3932 (1990).

31. E. S. Reckzeh, H. Waldmann, Small-Molecule Inhibition of Glucose Transporters GLUT-1-4. Chembiochem 21, 45–52 (2020).

32. B. Sun, H. Chen, J. Xue, P. Li, X. Fu, The role of GLUT2 in glucose metabolism in multiple organs and tissues. Mol Biol Rep 50, 6963–6974 (2023).

33. P. Iozzo et al., Insulin stimulates liver glucose uptake in humans: an 18F-FDG PET Study. J Nucl Med 44, 682–689 (2003).

34. M. C. Moore, K. C. Coate, J. J. Winnick, Z. An, A. D. Cherrington, Regulation of hepatic glucose uptake and storage in vivo. Adv Nutr 3, 286–294 (2012).

35. P. Thomas, Y. Pang, J. Dong, Enhancement of cell surface expression and receptor functions of membrane progestin receptor alpha (mPRalpha) by progesterone receptor membrane component 1 (PGRMC1): evidence for a role of PGRMC1 as an adaptor protein for steroid receptors. Endocrinology 155, 1107–1119 (2014).

36. Y. Pang, J. Dong, P. Thomas, Progesterone increases nitric oxide synthesis in human vascular endothelial cells through activation of membrane progesterone receptor-alpha. Am J Physiol Endocrinol Metab 308, E899–911 (2015).

37. J. C. Gonzalez-Orozco et al., Activation of membrane progesterone receptor-alpha increases proliferation, migration, and invasion of human glioblastoma cells. Mol Cell Endocrinol 477, 81–89 (2018).

38. W. Tan, J. Aizen, P. Thomas, Membrane progestin receptor alpha mediates progestin-induced sperm hypermotility and increased fertilization success in southern flounder (Paralichthys lethostigma). Gen Comp Endocrinol 200, 18–26 (2014).

39. W. Tan, P. Thomas, Activation of the Pi3k/Akt pathway and modulation of phosphodiesterase activity via membrane progestin receptor-alpha (mPRalpha) regulate progestin-initiated sperm hypermotility in Atlantic croaker. Biol Reprod 90, 105 (2014).

40. W. Tan, P. Thomas, Involvement of epidermal growth factor receptors and mitogen-activated protein kinase in progestin-induction of sperm hypermotility in Atlantic croaker through membrane progestin receptor-alpha. Mol Cell Endocrinol 414, 194–201 (2015).

41. E. Cabrera-Munoz, O. T. Hernandez-Hernandez, I. Camacho-Arroyo, Role of progesterone in human astrocytomas growth. Curr Top Med Chem 11, 1663–1667 (2011).

42. B. T. Hennessy, R. L. Coleman, M. Markman, Ovarian cancer. Lancet 374, 1371–1382 (2009).

43. C. A. Lange, D. Yee, Progesterone and breast cancer. Womens Health (Lond) 4, 151–162 (2008).

44. C. H. Diep, A. R. Daniel, L. J. Mauro, T. P. Knutson, C. A. Lange, Progesterone action in breast, uterine, and ovarian cancers. J Mol Endocrinol 54, R31–53 (2015).

45. P. Thomas, Membrane Progesterone Receptors (mPRs, PAQRs): Review of Structural and Signaling Characteristics. Cells 11, (2022).

46. J. Kim, G. Yang, Y. Kim, J. Kim, J. Ha, AMPK activators: mechanisms of action and physiological activities. Exp Mol Med 48, e224 (2016).

47. S. Herzig, R. J. Shaw, AMPK: guardian of metabolism and mitochondrial homeostasis. Nat Rev Mol Cell Biol 19, 121–135 (2018).

48. L. L. V. Moller, A. Klip, L. Sylow, Rho GTPases-Emerging Regulators of Glucose Homeostasis and Metabolic Health. Cells 8, (2019).

49. G. Leprivier, B. Rotblat, How does mTOR sense glucose starvation? AMPK is the usual suspect. Cell Death Discov 6, 27 (2020).

50. K. Inoki, J. Kim, K. L. Guan, AMPK and mTOR in cellular energy homeostasis and drug targets. Annu Rev Pharmacol Toxicol 52, 381–400 (2012).

51. K. G. de la Cruz López, M. E. Toledo Guzmán, E. O. Sánchez, A. García Carrancá, mTORC1 as a Regulator of Mitochondrial Functions and a Therapeutic Target in Cancer. Front Oncol 9, 1373 (2019).

52. S. H. Um, D. D’Alessio, G. Thomas, Nutrient overload, insulin resistance, and ribosomal protein S6 kinase 1, S6K1. Cell Metab 3, 393–402 (2006).

53. F. Tremblay et al., Identification of IRS-1 Ser-1101 as a target of S6K1 in nutrient- and obesity-induced insulin resistance. Proc Natl Acad Sci U S A 104, 14056–14061 (2007).

54. B. Viollet et al., Activation of AMP-activated protein kinase in the liver: a new strategy for the management of metabolic hepatic disorders. J Physiol 574, 41–53 (2006).

55. D. G. Hardie, F. A. Ross, S. A. Hawley, AMPK: a nutrient and energy sensor that maintains energy homeostasis. Nat Rev Mol Cell Biol 13, 251–262 (2012).

56. G. Zhou et al., Role of AMP-activated protein kinase in mechanism of metformin action. J. Clin. Invest. 108, 1167–1174 (2001).

57. Y. J. Choi et al., Inhibitory effect of mTOR activator MHY1485 on autophagy: suppression of lysosomal fusion. PLoS One 7, e43418 (2012).

58. T. L. van Belle, K. T. Coppieters, M. G. von Herrath, Type 1 diabetes: etiology, immunology, and therapeutic strategies. Physiol. Rev. 91, 79–118 (2011).

59. B. R. Miller, H. Nguyen, C. J. Hu, C. Lin, Q. T. Nguyen, New and emerging drugs and targets for type 2 diabetes: reviewing the evidence. Am Health Drug Benefits 7, 452–463 (2014).

60. S. D. Pastakia, C. R. Pekny, S. M. Manyara, L. Fischer, Diabetes in sub-Saharan Africa - from policy to practice to progress: targeting the existing gaps for future care for diabetes. Diabetes Metab Syndr Obes 10, 247–263 (2017).

61. R. A. DeFronzo et al., Type 2 diabetes mellitus. Nat Rev Dis Primers 1, 15019 (2015).

62. J. B. Buse et al., 2019 Update to: Management of Hyperglycemia in Type 2 Diabetes, 2018. A Consensus Report by the American Diabetes Association (ADA) and the European Association for the Study of Diabetes (EASD). Diabetes Care 43, 487–493 (2020).

63. J. P. H. Wilding et al., Once-Weekly Semaglutide in Adults with Overweight or Obesity. N Engl J Med 384, 989–1002 (2021).

64. D. Moriconi et al., Predictors of type 2 diabetes relapse after Roux-en-Y Gastric Bypass: A ten-year follow-up study. Diabetes Metab 48, 101282 (2022).

65. M. Foretz, B. Guigas, B. Viollet, Metformin: update on mechanisms of action and repurposing potential. Nat Rev Endocrinol 19, 460–476 (2023).

66. G. R. Steinberg, D. Carling, AMP-activated protein kinase: the current landscape for drug development. Nat Rev Drug Discov 18, 527–551 (2019).

67. W. W. Wheaton et al., Metformin inhibits mitochondrial complex I of cancer cells to reduce tumorigenesis. Elife 3, e02242 (2014).

68. E. Fontaine, Metformin-Induced Mitochondrial Complex I Inhibition: Facts, Uncertainties, and Consequences. Front Endocrinol (Lausanne) 9, 753 (2018).

69. F. Bonnet, A. Scheen, Understanding and overcoming metformin gastrointestinal intolerance. Diabetes Obes Metab 19, 473–481 (2017).

70. G. E. Dressing, R. Alyea, Y. Pang, P. Thomas, Membrane progesterone receptors (mPRs) mediate progestin induced antimorbidity in breast cancer cells and are expressed in human breast tumors. Horm. Cancer 3, 101–112 (2012).

71. L. Zuo, W. Li, S. You, Progesterone reverses the mesenchymal phenotypes of basal phenotype breast cancer cells via a membrane progesterone receptor mediated pathway. Breast Cancer Res 12, R34 (2010).

72. S. Chen et al., PAQR8 promotes breast cancer recurrence and confers resistance to multiple therapies. Breast Cancer Res 25, 1 (2023).

73. M. Sinreih, T. Knific, P. Thomas, S. Frkovic Grazio, T. L. Rizner, Membrane progesterone receptors beta and gamma have potential as prognostic biomarkers of endometrial cancer. J. Steroid Biochem. Mol. Biol. 178, 303–311 (2018).

74. B. Li, Z. Lin, Q. Liang, Y. Hu, W. F. Xu, PAQR6 Expression Enhancement Suggests a Worse Prognosis in Prostate Cancer Patients. Open Life Sci 13, 511–517 (2018).

